# Why concatenation fails in the anomaly zone

**DOI:** 10.1101/116509

**Authors:** Fábio K. Mendes, Matthew W. Hahn

## Abstract

Genome-scale sequencing has been of great benefit in recovering species trees, but has not provided final answers. Despite the rapid accumulation of molecular sequences, resolving short and deep branches of the tree of life has remained a challenge, and has prompted the development of new strategies that can make the best use of available data. One such strategy – the concatenation of gene alignments – can be successful when coupled with many tree estimation methods, but has also been shown to fail when there are high levels of incomplete lineage sorting. Here, we focus on the failure of likelihood-based methods in retrieving a rooted, asymmetric four-taxon species tree from concatenated data when the species tree is in or near the anomaly zone – a region of parameter space where the most common gene tree does not match the species tree because of incomplete lineage sorting. First, we use coalescent theory to prove that most informative sites will support the species tree in the anomaly zone, and that as a consequence maximum-parsimony succeeds in recovering the species tree from concatenated data. We further show that maximum-likelihood tree estimation from concatenated data fails both inside and outside the anomaly zone, and that this failure is unconnected to the frequency of the most common gene tree. We provide support for a hypothesis that likelihood-based methods fail in and near the anomaly zone because discordant sites on the species tree have a lower likelihood than those that are discordant on alternative topologies. Our results confirm and extend previous reports of the failure and success of likelihood- and parsimony-based methods, and highlight avenues for future work improving the performance of methods aimed at recovering species tree.

---

One of the major goals of evolutionary biology is the reconstruction of species relationships (Edwards 2009). Species trees – or phylogenies – are valued end products themselves (Hinchliff et al. 2015), but it is perhaps their central role in comparative studies that makes their accurate reconstruction so critical. Comparative analyses can include inferences about trait evolution, the dynamics of extinction and speciation, and species divergence times (O’Meara 2012; Hahn and Nakhleh 2016).

Unfortunately, the quest of reconstructing phylogenies has always been a difficult one. A scarcity of data was a major hurdle in phylogenetic analyses for 30 years after the birth of molecular systematics (Zuckerkandl and Pauling 1965) due to the technical and financial challenges of DNA and protein sequencing. Small datasets meant that sampling error was likely to occur, and the resulting disagreement between inferred trees had to be reconciled (Slowinski and Page 1999). As a consequence, a major topic of contention was whether and how to combine datasets (Huelsenbeck et al. 1996; Page 1996).

With the accumulation of larger datasets, a practice that became known as “concatenation” was widely adopted (Philippe et al. 2005; Edwards 2009). Concatenation is an intuitive procedure aimed at combining the information contained in the sequences of many genes in a single analysis. Concatenated datasets were easily analyzed by existing tree-building methods, including those employing explicit models of sequence evolution (which had come to dominate phylogenetics by then; Steel and Penny 2000). Initial studies using concatenation yielded high-confidence phylogenies from genes whose individual trees were often discordant (e.g., Soltis et al. 1999; Murphy et al. 2001; Rokas et al. 2003). Concatenation was therefore seen as holding the promise to end the problem of sampling error and the incongruence it produced (Gee 2003).

The amassing of more genes – followed by concatenation – is indeed expected to reduce the amount of noise due to sampling error. Many phenomena, however, pose difficulties to this approach because they produce discordant trees for biological reasons. Among these phenomena, incomplete lineage sorting (ILS) is perhaps the most well studied, partly because it is conducive to modeling and mathematical characterization (Hudson 1983; Tajima 1983; Pamilo and Nei 1988). Going backwards in time, ILS is said to occur when lineages from the same population fail to coalesce, and instead coalesce in an ancestral population. As a result, they may coalesce with more distantly related lineages, leading to discordance. ILS is relevant to all phylogenetic analyses because it results from an inherent property of natural populations, and has accordingly been shown to be pervasive across the tree of life (e.g., Pollard et al. 2006; White et al. 2009; Hobolth et al. 2011; Brawand et al. 2014; Zhang et al. 2014; Suh et al. 2015; Pease et al. 2016). Combining loci that are discordant due to ILS means that concatenation analyses will be averaging over many different topologies; the hope is that the most common pattern will coincide with the true species relationships.

However, sometimes the most common gene tree topology does not coincide with the species tree: in extreme cases ILS can produce unexpected results in an area of tree space called the “anomaly zone” (AZ; Degnan and Rosenberg 2006). ILS is increasingly more likely as species tree internal branches get shorter (i.e., as the time between two or more speciation events is shorter). When two or more consecutive internal branches on a species tree are sufficiently short, gene trees incongruent with the species tree can be more common than congruent gene trees (Fig. 1; Degnan and Salter 2005; Degnan and Rosenberg 2006). In other words, inside the AZ the topology of the most common gene tree (also referred to as the “anomalous gene tree” [AGT]; Fig. 1) in the dataset does not match that of the species tree. The AZ was found to be particularly troublesome for species tree estimation, as it was shown via simulation that for species trees inside the AZ, concatenation can lead to a maximum-likelihood tree whose topology matches the AGT rather than the species tree (Kubatko and Degnan 2007). The finding that the AZ could pose problems for species tree estimation motivated a decade’s worth of research into methods that could recover the correct species relationships (e.g., Liu and Pearl 2007; Liu et al. 2009, 2010; Heled and Drummond 2010; Larget et al. 2010; Mirarab and Warnow 2015). Along with these new methods came studies into the behavior of traditional tree inference methods on concatenated data in the AZ (e.g., Liu and Edwards 2009) and the conditions under which the AZ could impact empirical studies (e.g., Huang and Knowles 2009).

**Figure 1:**
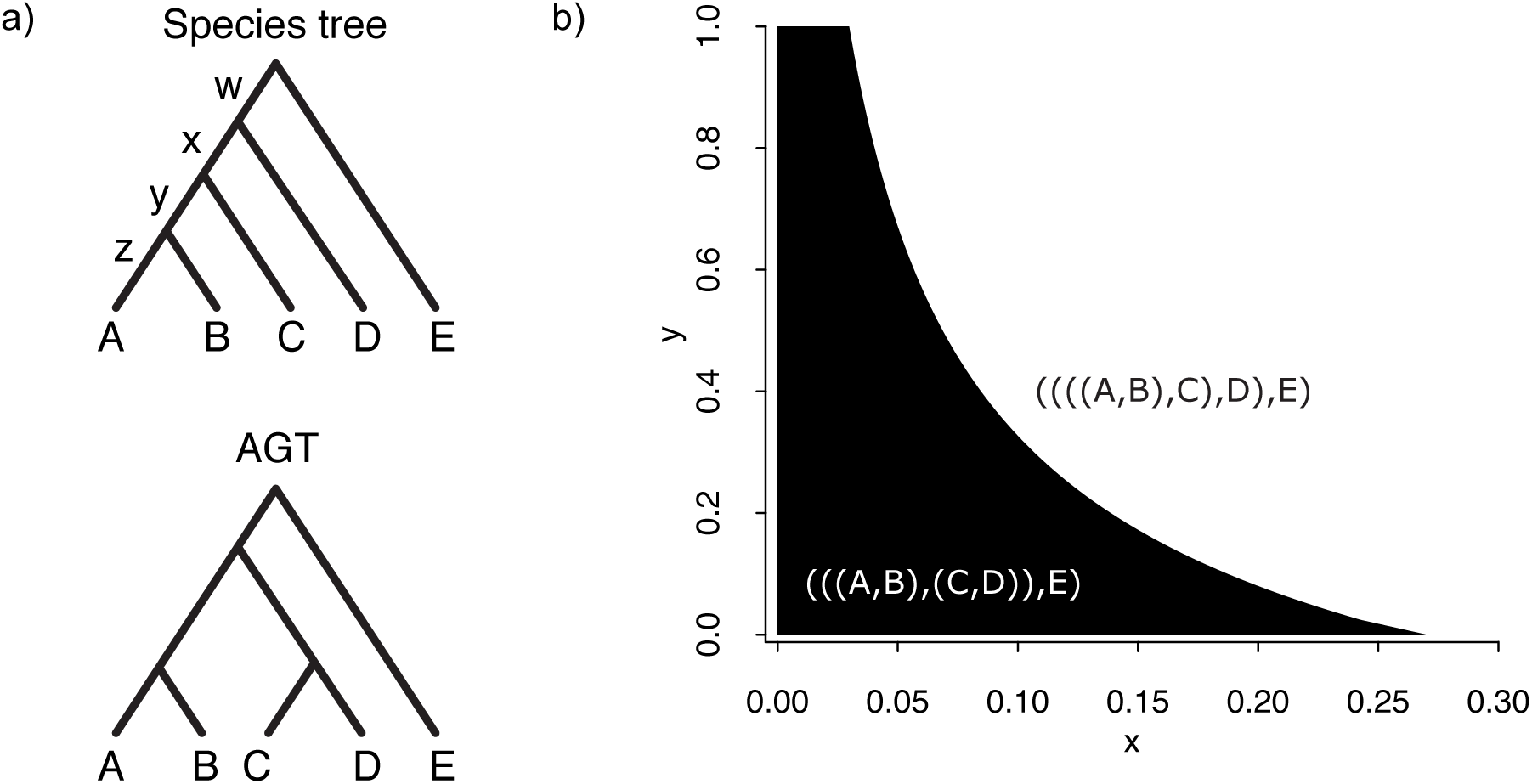
(a) Top tree: smallest species tree for which an anomaly zone can be defined, where *z* is the length of terminal branches A and B, and *w*, *x* and *y* are the lengths of the three internal branches (oldest to youngest), respectively. Bottom tree: the most common gene tree (an anomalous gene tree, AGT) when species tree ((((A,B),C),D),E) (top tree) is inside the anomaly zone. Branch lengths are arbitrary and were not drawn in proportion to theoretical or simulated averages. (b) Phylogenetic tree space for species tree ((((A,B),C),D),E), where *x* and *y* correspond to the lengths of the oldest and youngest ingroup internal branches, respectively (as shown in [a]; *x* and *y* are measured in coalescent units, i.e., *N*_e_ generations). The region shaded in black corresponds to the anomaly zone (Degnan and Rosenberg 2006), in which the most common gene tree is AGT (((A,B),(C,D)),E).

Here, we explore two interesting results from this literature, and their implications for phylogenetic reconstruction: (i) Both parsimony- and distance-based methods appear to succeed in inferring the species tree *inside* the AZ (Liu and Edwards 2009), and (ii) Maximum-likelihood species tree estimates from concatenated data can be incorrect just *outside* the AZ (Kubatko and Degnan 2007). These two observations are not consistent with the accepted narrative concerning problems with concatenation inside the AZ, but both appear to be correct (see below). To explore these results – and the behavior of concatenation more generally – we use coalescent theory to mathematically demonstrate why parsimony succeeds inside the AZ for a rooted species tree with four taxa. We then provide an explanation as to why maximum-likelihood applied to concatenation can fail both inside and outside the AZ. In fact, we show that the failure of such approaches is not directly tied to the AZ at all. Our results cast doubt on the seemingly common notion that concatenation is the sole culprit to blame for incorrect species tree estimates from datasets in the AZ. Finally, we suggest future research directions in light of our results.

## Most informative sites in the anomaly zone support the species tree

Theoretical results concerning the anomaly zone have focused on the distribution and frequencies of different gene trees (Degnan and Rosenberg 2006, 2009; Rosenberg and Tao 2008). If species trees are constructed by simply taking the most common gene tree (also known as “democratic vote”), then in the AZ the species tree will incorrectly be inferred to be the AGT. However, this method is rarely used, and is not directly relevant to concatenated analyses unless the most common site pattern also supports the most common gene tree. Informative site patterns in molecular phylogenetics are the result of substitutions occurring along the internal branches of a gene tree (here we use the term “gene” to mean any non-recombining genomic segment). Despite the greater frequency of anomalous gene trees in the AZ (compared to congruent gene trees), because of their very short internal branches we hypothesized that the most common site patterns would still support the species tree. If this is the case, parsimony methods would support the species tree inside the AZ.

In order for site patterns supporting the species tree to be the most common, the total length of concordant internal branches (i.e., the sum of lengths, over all gene trees, of internal branches that exist in the species tree) must be greater than that of internal branches supporting any of the other unique topologies. Under an infinite-sites mutation model the topology supported by the greatest total internal branch length will also be supported by the largest number of informative site patterns. Therefore, to determine the expected number of informative site patterns supporting any topology when a large number of gene trees are sampled, one must know (i) the probability of all gene tree topologies, and (ii) the expected lengths of the internal branches present in each of these topologies. Knowing (i) and (ii) allows the calculation of *S_t_,* the total length of internal branches supporting any topology *t* in *T,* the set of all possible gene tree topologies under the species tree. In the case of a rooted four-taxon species tree, for example, there are 15 possible topologies, and so |*T*| = 15 with *t* taking any value from 1 to 15 (Table 1). *S_t_* can then be computed for any of these 15 topologies.

**Table 1:** Probability of each gene tree topology under species tree ((((A,B),C),D),E) (where E is the outgroup), when ILS is the sole cause of incongruence (from Table V and Equation 2 in Rosenberg, 2002). Branches *y* and x are the most recent and oldest ingroup internal branches, respectively, with lengths expressed in coalescent units.

| Topology | $t$ or $u$ | $P(u)$ |
| --- | --- | --- |
| ((A,B),(C,D)) | 1 | $\frac{1}{3}e^{-x} - \frac{1}{6}e^{-(x+y)} - \frac{1}{18}e^{-(3x+y)}$ |
| ((A,C),(B,D)) | 2 | $\frac{1}{6}e^{-(x+y)} - \frac{1}{18}e^{-(3x+y)}$ |
| ((B,C),(A,D)) | 3 | $\frac{1}{6}e^{-(x+y)} - \frac{1}{18}e^{-(3x+y)}$ |
| (((A,B),C),D) | 4 | $1 - \frac{2}{3}e^{-x} - \frac{2}{3}e^{-y} + \frac{1}{3}e^{-(x+y)} + \frac{1}{18}e^{-(3x+y)}$ |
| (((A,B),D),C) | 5 | $\frac{1}{3}e^{-x} - \frac{1}{6}e^{-(x+y)} - \frac{1}{9}e^{-(3x+y)}$ |
| (((A,C),B),D) | 6 | $\frac{1}{3}e^{-x} - \frac{1}{6}e^{-(x+y)} + \frac{1}{18}e^{-(3x+y)}$ |
| (((A,C),D),B) | 7 | $\frac{1}{6}e^{-(x+y)} - \frac{1}{9}e^{-(3x+y)}$ |
| (((A,D),B),C) | 8 | $\frac{1}{18}e^{-(3x+y)}$ |
| (((A,D),C),B) | 9 | $\frac{1}{18}e^{-(3x+y)}$ |
| (((B,C),A),D) | 10 | $\frac{1}{3}e^{-x} - \frac{1}{6}e^{-(x+y)} + \frac{1}{18}e^{-(3x+y)}$ |
| (((B,C),D),A) | 11 | $\frac{1}{6}e^{-(x+y)} - \frac{1}{9}e^{-(3x+y)}$ |
| (((B,D),A),C) | 12 | $\frac{1}{18}e^{-(3x+y)}$ |
| (((B,D),C),A) | 13 | $\frac{1}{18}e^{-(3x+y)}$ |
| (((C,D),A),B) | 14 | $\frac{1}{18}e^{-(3x+y)}$ |
| (((C,D),B),A) | 15 | $\frac{1}{18}e^{-(3x+y)}$ |

Computing *S_t_* is done by first identifying the set of all topologies, *U,* sharing internal branches with *t*, and recording the probability of each topology, *u*, in *U.* We denote these probabilities *P*(*u*). Second, for each topology *u*, we must identify the set of all internal branches, *B_u_,_t_,* that it shares with *t.* Each branch *b* in *B_u_,_t_* is labeled with a number from 1 to |*B_u,t_*|, and we record the expected length of each branch *b* given *u,* L(*b*|*u*). Therefore, *S_t_* can be calculated as:

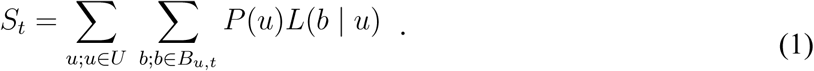

Whichever topology *t* maximizes *S_t_* will by definition be supported by the largest number of informative site patterns, and will usually also be the most parsimonious tree.

When ILS is the only cause of phylogenetic incongruence, both the probability of observing each different gene tree topology and the expected lengths of their internal branches will be functions of the species tree’s internal branch lengths. In the case of the four-taxon species tree ((((A,B),C),D),E) (where E is the outgroup, henceforth omitted from parenthetic notation), the probability of each of the 15 possible topologies has been derived (Table 1; Rosenberg 2002). Under this species tree, the most common gene tree will always be either (((A,B),C),D) (outside the AZ; Fig. 1b) or ((A,B),(C,D)) (inside the AZ; Fig. 1b). In evaluating the strength of support for the species tree versus the AGT, we can simplify our calculations by noting that these two competing topologies differ in only the single, deepest internal branch: this branch subtends ((A,B),C) in the congruent gene tree, while in the AGT it subtends (C,D) (the internal branch leading to (A,B) is shared by both topologies; Fig. 1a). Therefore, understanding which topology is supported by the most informative site patterns inside the AZ only requires us to compare the total length of branches subtending ((A,B),C) to that of branches subtending (C,D).

A closer look at the 15 distinct gene tree topologies reveals that only six are relevant to these two internal branches (Fig. 2). The topologies (((A,B),C),D), (((A,C),B),D) and (((B,C),A),D) (*u* = 4, 6 and 10, respectively; Table 1) share one internal branch each with the species tree topology (i.e., |*B_4_,_4_*| = |*B_6_,_4_*| = |*B_10_,_4_*| = 1; Fig. 2), while the topologies ((A,B),(C,D)), (((C,D),A),B), and (((C,D),B),A) (*u* = 1, 14 and 15, respectively; Table 1) share one internal branch each with the AGT (Fig. 2). Coalescent theory can be used to find the expected frequency of these topologies and the length of the relevant branches within them.

**Figure 2:**
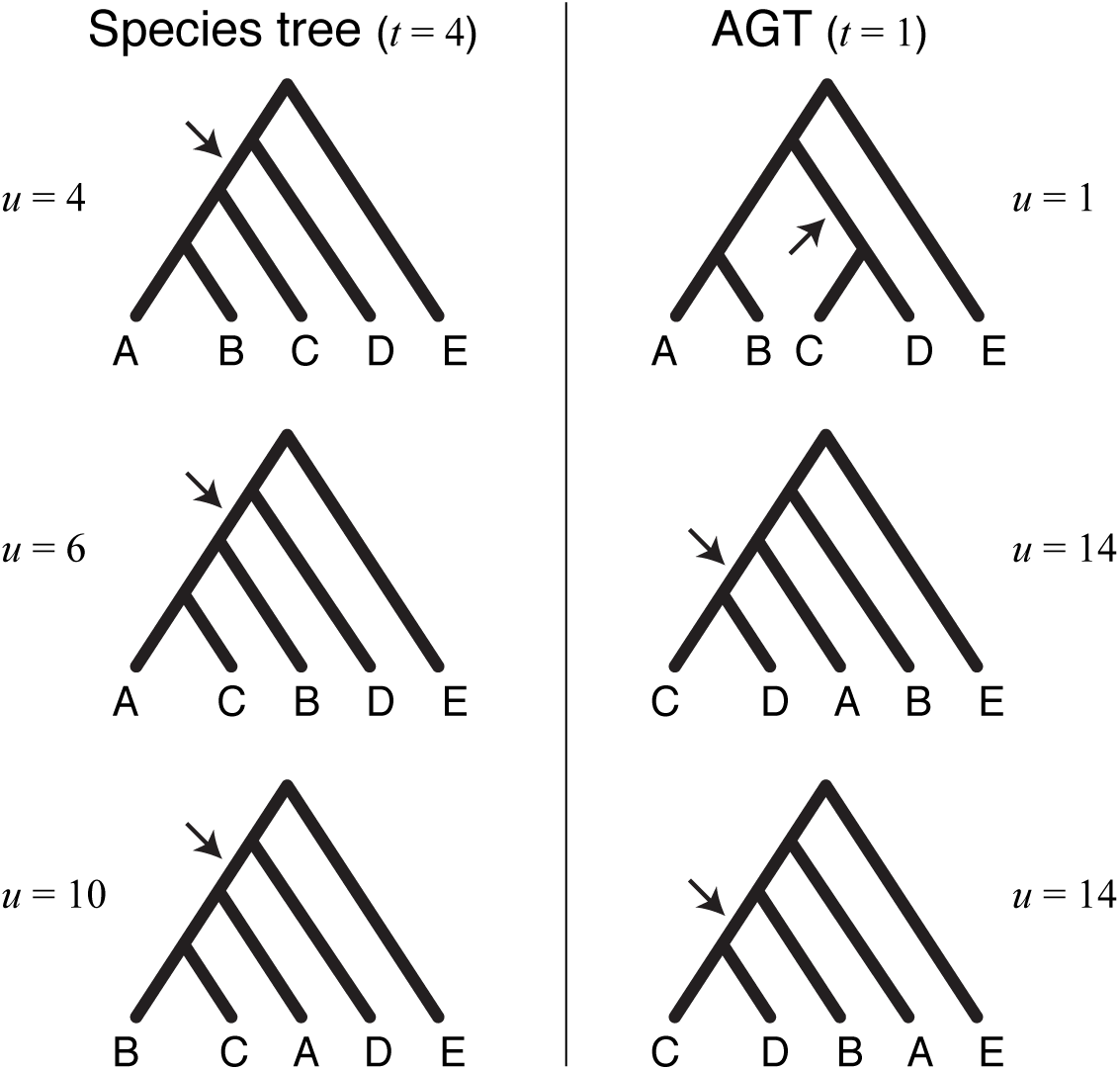
Gene tree topologies that give support to the species tree (left column; *t* = 4, Table 1) or to the AGT (right column; *t* = 1, Table 1). Arrows indicate the internal branch that each gene tree topology *u* contributes to topology *t*. Branch lengths are arbitrary and were not drawn in proportion to theoretical or simulated averages.

For species tree (((A,B),C),D), application of equation 1 shows that the AGT ((A,B),(C,D)) should never have more sites supporting it than the species tree. Even in the most extreme scenario, when internal branch lengths *x* and *y* are zero, the species tree (SP) and AGT ((A,B),(C,D)) are equally supported:

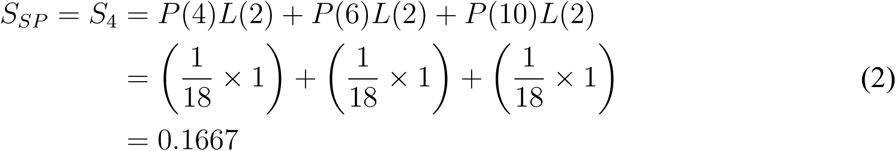

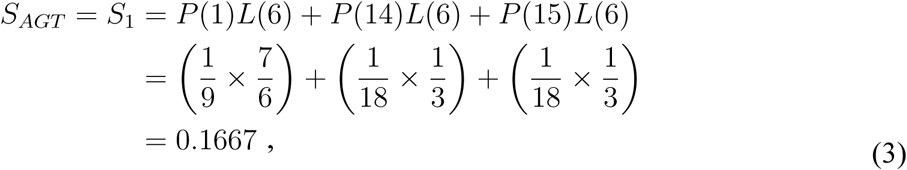

where branch lengths are given in coalescent units, and for each *u* the single branch being considered is labeled *b* = 1 (Fig. 2).

Because the probability of observing the congruent gene tree only increases as the x and *y* branch lengths in the species tree become larger, the total length of internal branches over all gene trees supporting the species tree will always be greater than that supporting the AGT. We derive expected values for any *x* and *y* in Appendix A. As a result, even though the most common tree matches the AGT, the most common site pattern supports the species tree. Therefore, parsimony-based methods should accurately recover the species tree topology in the AZ.

In order to confirm our theoretical expectations, we performed coalescent simulations across parameter space for species tree (((A,B),C),D). More specifically, we simulated 20,000 gene trees at each of multiple coordinates forming a grid across tree space (Fig. 3; see details in Appendix B). First, we recorded the most common gene tree at each coordinate and observed a very close match with the theoretical AZ (Degnan and Rosenberg 2006; Supplementary Fig. 1). We then simulated one 1-kb nucleotide sequence per gene tree using the Jukes-Cantor model (Jukes and Cantor 1969), and concatenated all 20,000 sequences into one single alignment per grid coordinate. By using maximum parsimony to estimate the species tree from each concatenated alignment we were able to recapitulate Liu and Edwards' (2009) result: the estimated species tree species was congruent with the true species tree across all of tree space (Fig. 3; the same was true using neighbor-joining on the concatenated alignments; result not shown). Finally, we compared the expected “SP:AGT” ratio of the total lengths of internal branches supporting either topology (i.e., *S_4_:S_1_;* see Equation 1) to the simulated ratio (obtained by summing simulated gene tree internal branch lengths). This was done for 19 different pairs of *x* and *y* values, and for each pair we replicated our simulations 100 times (each replicate consisted of 20,000 simulated gene trees). The expected ratio was closely approximated by the simulated ratio (Fig. 4).

**Figure 3:**
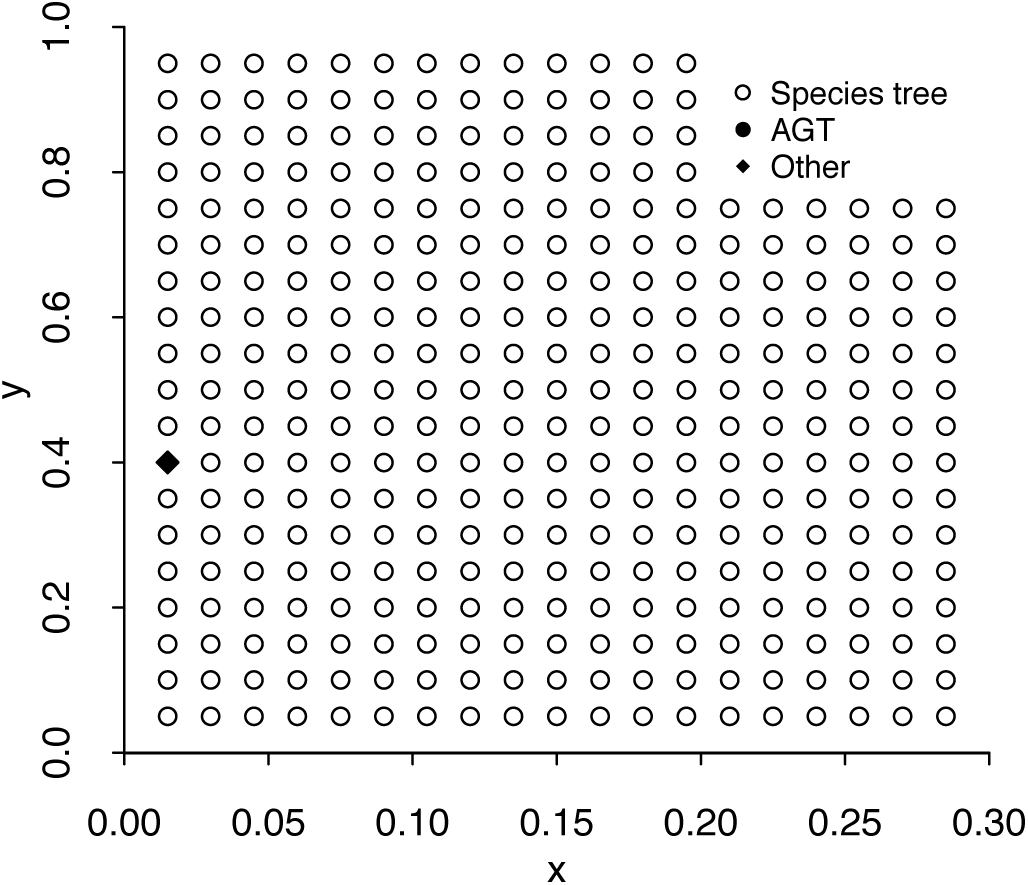
Topology reconstructed by parsimony across the tree space of species tree ((((A,B),C),D),E). The phylogeny at each grid point was estimated from a concatenated alignment of 20,000 1-kb loci generated under the multispecies coalescent simulated at that coordinate of tree space. Branches *x* and *y* are measured in coalescent units.

**Figure 4:**
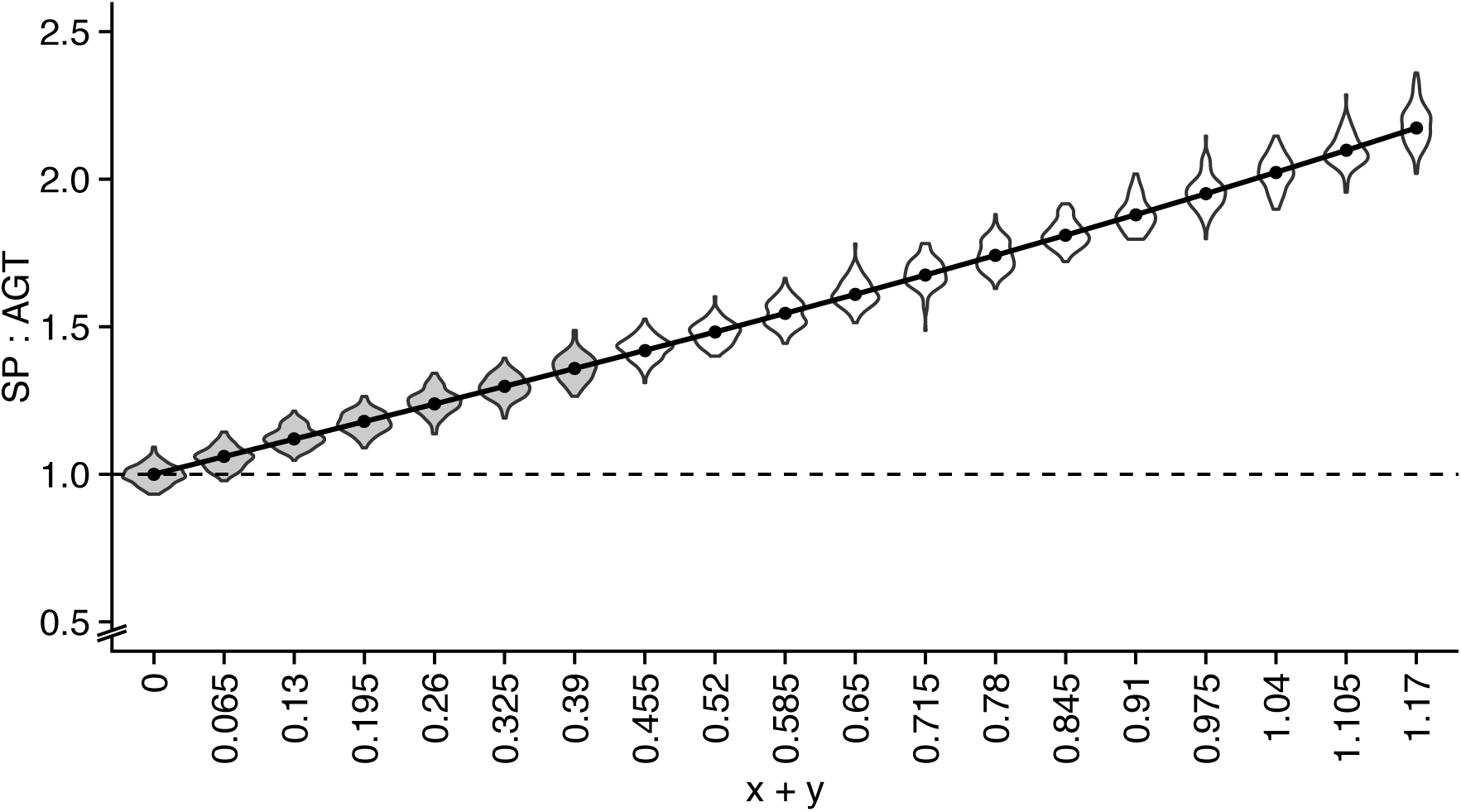
Expected (connected dots) and simulated (100 replicates per coordinate; violin plots) support for species tree ((((A,B),C),D),E) and AGT (((A,B),(C,D)),E) expressed as a ratio of total internal branch lengths supporting either topology, at 19 coordinates across tree space (see Appendix B). Coordinates for which violin plots are shaded in gray are located inside the anomaly zone. The sum of x and *y* is in coalescent units.

Our results suggest that parsimony succeeds inside the AZ for a four-taxon rooted tree because there will always be more sites supporting the species tree topology than any other topology. This is in contrast to the explanation put forward by Liu and Edwards (2009) for why parsimony correctly recovers the species tree in the AZ. In their simulations, the site pattern supporting the species tree was also observed to be the most common, but this outcome was interpreted to be a result of long-branch attraction (LBA; (Felsenstein 1978) biasing parsimony against the AGT. They concluded that parsimony was therefore getting the right answer for the wrong reasons (Liu and Edwards 2009). We note that given the value of *θ* (the population mutation parameter) used in our simulations, terminal branches are not close to being saturated, and so LBA is not biasing parsimony against the AGT.

When concatenating sequences, clarifying the distinction between the most common gene tree and the most common site pattern in the dataset is critical: even if the most common gene tree is incongruent, more site patterns can still support the congruent gene tree because they come from multiple different topologies each with longer internal branches on average. For the asymmetric species tree with four taxa, we should always expect more site patterns supporting the species tree rather than the AGT (Fig. 4). Therefore, when parsimony- and distance-based methods succeed in reconstructing the species tree, they both do so for the *right* reasons.

Hence for the species tree considered here, concatenation cannot be causing phylogenetic reconstruction methods to fail *per se.* Concatenation is expected to remove sampling noise, and as long as there is more phylogenetic signal supporting the species tree than supporting any other topology, concatenation should not interfere with species tree reconstruction when using counts of informative site patterns (i.e., parsimony). Because the phylogenetic signal supporting the species tree topology (((A,B),C),D) is always higher than that supporting AGT ((A,B),(C,D)), our results imply that when species tree reconstruction from concatenated datasets fail, it must be due to properties of likelihood-based methods, not concatenation.

## Species tree reconstruction from concatenated data fails because of likelihood

### The anomaly zone is not directly connected to the failure of likelihood methods

In the previous section we showed that parsimony-based methods are expected to succeed for species tree (((A,B),C),D) in all areas of tree space examined. There are always more sites supporting the species tree than the most common gene tree in the dataset, which is the AGT [((A,B),(C,D))]. These results have two implications. First, as mentioned above, concatenation *per se* is not responsible for the failure of tree reconstruction in the AZ. It must be that likelihood-based methods fail because of properties of these methods when applied to concatenated datasets. Second, the above results suggest that the region in tree space where likelihood-based methods fail does not necessarily coincide with the AZ. If such methods are failing for reasons other than the frequency of the most common gene tree, then there is no reason that their failure should follow the frequency of the most common gene tree (i.e., the AZ). This second implication is supported by our results and by previous observations that likelihood-based methods can fail even *outside* the AZ, and succeed *inside* the AZ (Kubatko and Degnan 2007).

Similar to what was done in our investigation of parsimony- and distance-based methods described in the previous section, we examined the performance of maximum-likelihood estimation across the tree space of species tree (((A,B),C),D). We used the same concatenated alignments at the same coordinates of tree space, and recorded the maximum-likelihood tree at each grid point (see details in Appendix B). In agreement with many previous studies (e.g., Kubatko and Degnan 2007; Liu and Edwards 2009), species tree reconstruction with maximum-likelihood on concatenated data failed at many points in the AZ (Fig. 5).

**Figure 5:**
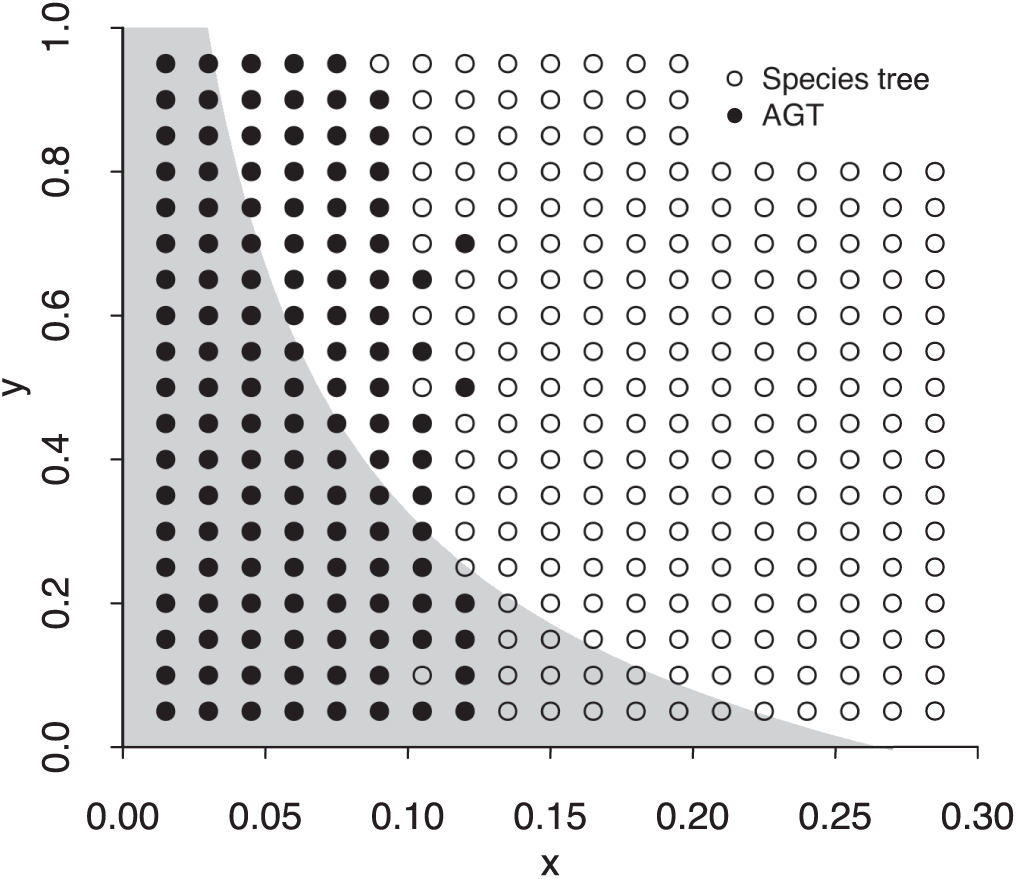
Topology reconstructed using maximum-likelihood across the tree space of species tree ((((A,B),C),D),E). The phylogeny at each grid point was estimated from a concatenated alignment of 20,000 1-kb loci generated under the multispecies coalescent simulated at that coordinate of tree space (these are the same alignments used in Figure 3). Branches *x* and *y* are measured in coalescent units. The region shaded in gray corresponds to the anomaly zone (Degnan and Rosenberg 2006), in which the most common gene tree is the AGT.

However, because we covered parameter space more extensively than previous investigations, we are able to observe clear regions inside the AZ where maximum-likelihood succeeds in recovering the species tree, instead of just a few coordinates in tree space near the AZ border (Fig. 5). We also identified a region outside the AZ where the AGT was favored by maximum-likelihood (Fig. 5). Our results confirm the disconnection between the AZ and the area of parameter space in which likelihood-based methods applied to concatenated data seem to be inconsistent.

These results support the conclusions drawn from the previous section: the failure of analyses using concatenation is not due to the identity of the most frequent gene tree topology. But these observations beg the questions of why the maximum-likelihood tree differs from the most parsimonious tree, and what determines the shape of the region in tree space in which maximum-likelihood estimation seems to be inconsistent. We address these questions in the next section.

### The cost of discordant sites explains why likelihood-based methods fail to reconstruct the species tree from concatenated data

While likelihood-based methods have many advantages over other classes of methods (e.g., Huelsenbeck 1995; Swofford et al. 2001; Ogden and Rosenberg 2006), the demonstration that maximum-likelihood tree estimation can fail to reconstruct the species tree is not entirely surprising. When models are mis-specified, likelihood-based methods can be unsuccessful in recovering the true tree, and in such cases these methods have been shown to converge on the wrong answer (e.g., Gaut and Lewis 1995; Sullivan and Swofford 1997). Although maximum-likelihood estimation from concatenated data can be robust to low ILS levels (Tonini et al. 2015; Mirarab et al. 2016), the success of this approach is clearly not guaranteed when there are high levels of ILS. So what is the nature of model inadequacy when concatenated data is used in a maximum-likelihood framework? One possible answer is that, in cases involving ILS, concatenation violates the assumption that all sites have evolved along a single topology (Roch and Steel 2015). Because parsimony and neighbor-joining applied to concatenated datasets do not fail, however, the failure of maximum-likelihood estimation must be due to differences as to how discordant trees are accommodated by different methods. Below we offer one hypothesis to explain why likelihood favors the AGT over the species tree, but parsimony does not.

To better explain our hypothesis, it will be useful to first discuss one important consequence of including discordant tree topologies in an alignment. Relative to a focal tree, discordant topologies contain branches that do not exist in this tree (Robinson and Foulds 1981). For example, the AGT (tree on the right in Fig. 6) has an internal branch leading to the ancestor of C and D that is not present in the species tree. Conversely, if the AGT is our focal tree, then the species tree has an exclusive internal branch leading to the ancestor of A, B, and C. We refer to these as “discordant branches,” and to the site patterns produced by substitutions on them as “discordant sites” (all other sites are considered concordant). Such site patterns are particularly important because they must be resolved by proposing more than one substitution on the focal tree; we previously described this phenomenon, referring to the artefactual changes as “substitutions produced by ILS” (SPILS; Mendes and Hahn 2016). Site 1 in Fig. 6 shows an example of SPILS where a substitution occurring on a branch exclusive to the species tree would be inferred to have been due to two substitutions on the AGT. Site 2 shows the opposite pattern, as the substitution on the discordant branch of the AGT must be mapped twice onto the species tree.

**Figure 6:**
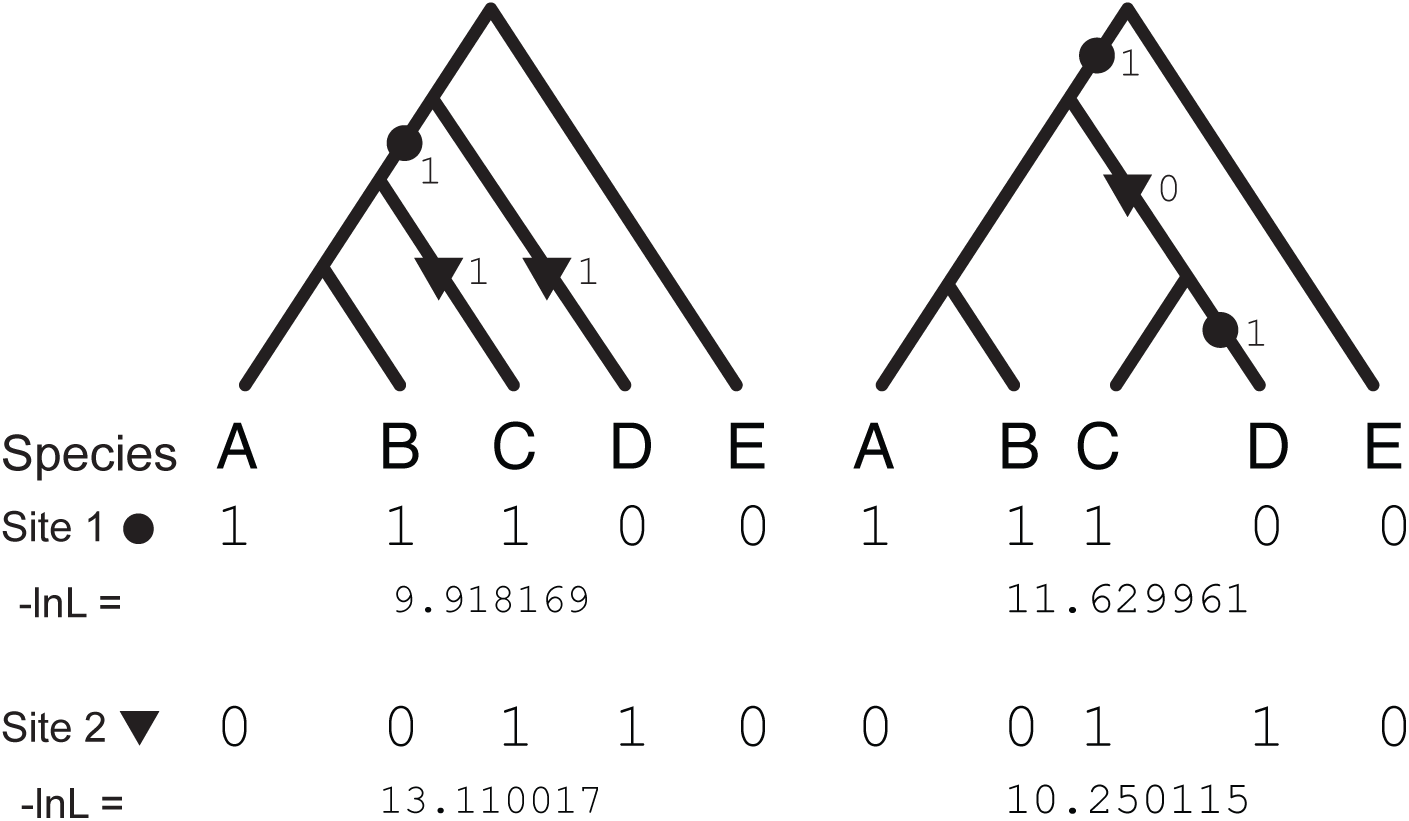
The species tree topology (left) and the anomalous gene tree topology (right). Filled circles and triangles represent character state transitions. Site 1 is concordant with the species tree and discordant with anomalous gene tree. Conversely, site 2 is concordant with the anomalous gene tree and discordant with the species tree. Negative log-likelihoods (-lnL) for each site pattern were computed on the maximum-likelihood tree obtained from concatenated data when *x* = 0.015 and *y* = 0.05.

In the presence of gene tree discordance, evaluating a tree on a concatenated alignment will thus entail considering both concordant and discordant site patterns. The key distinction between them is that discordant sites will always cost more on the focal tree than concordant sites. How the total score of a tree is calculated – and how parsimony and likelihood methods deal with these costs in particular – turns out to be crucial in understanding their behavior.

In the case of parsimony-based methods, if we define *O* and *D* as the sets of all possible concordant and discordant site patterns, respectively, the parsimony score of a tree *t*, *P_t_,* will be:

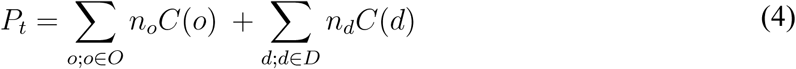

where *n_o_* and *C*(*o*) are the count and cost of concordant site pattern *o*, and *n_d_* and *C*(*d*) are the count and cost of discordant site pattern *d. C*(*o*) and *C*(*d*) equal the minimum number of substitutions required to generate site patterns *o* and *d*, respectively, on tree *t.* The tree *t* with the lowest parsimony score, *P_t_*, is considered the most parsimonious and will be preferred over less parsimonious ones.

Under a simple weighting scheme for a rooted four-taxon tree, it is easy to see that *C*(*o*) = 1 and *C*(*d*) = 2 for all possible (biallelic) site patterns: when a site pattern is concordant with a topology, it can always be resolved with a single substitution; when it is discordant, it can always be resolved with two substitutions (Fig. 6). The most parsimonious tree is therefore the one that maximizes *n_o_* (which directly reflects the value of *S_t_* in equation 1). For the rooted asymmetric four-taxon tree case, we have proven above that maximum-parsimony methods are consistent under this weighting scheme.

The likelihood score of a tree *t*, *L_t_,* can also be written in terms similar to those in equation 4:

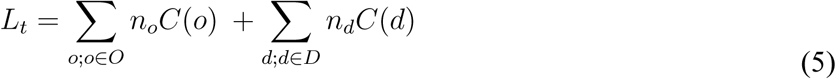

with the difference that *C*(*o*) and *C*(*d*) now correspond to the negative log-likelihoods of concordant site pattern *o*, and discordant site pattern *d*, respectively (the lower the negative log-likelihoods, the less costly and the more likely a site is). A crucial distinction is that in maximum-likelihood tree estimation, the likelihood that a site in the alignment contributes to the final tree likelihood depends on the mapping of nucleotide substitutions at all other sites, expressed as branch lengths. Branch lengths serve as a proxy for the expected probability of change, and so nucleotide transitions – including homoplasious ones – along longer branches are more probable (less “costly”) than along shorter ones. This property grants likelihood-based methods more robustness (compared to parsimony-based methods) to problems such as long-branch attraction, making them good choices for molecular phylogenetic analyses.

We hypothesize that it is precisely the attribute of likelihood-based methods mentioned above – their ability to take branch lengths into account when evaluating different trees – that contributes to their convergence on the incorrect tree topology in the presence of high levels of ILS. While the parsimony costs of concordant and discordant sites for the rooted four-taxon species tree are fixed at 1 and 2, respectively, this is not true for the likelihood costs. Negative log-likelihoods of concordant and discordant sites depend on the branch lengths of the trees on which they are evaluated. If the species tree and AGT differ in their branch lengths, so will the likelihood costs of concordant and discordant site patterns under either topology, leading to possibly different final tree likelihoods.

An immediately obvious difference in branch lengths between the two competing topologies considered here is the length of branches exclusive to each tree. Concordant site patterns for each topology will include substitutions occurring on these branches, and the costs, *C*(*o*), may therefore differ on the two topologies. A less obvious – but ultimately more important – difference is that sites that are differentially discordant on each topology (i.e., those sites that are not discordant on both) will be resolved along different branches that potentially have different lengths (e.g., site 1 in Fig. 6 is resolved along branches C and D of the AGT; site 2 is resolved on the species tree along branch D and the internal branch leading to the root). The difference in lengths between these two pairs of branches will then contribute to differences in final tree likelihoods. Therefore, unlike the case of maximum-parsimony described above, the maximum-likelihood tree is not simply the one for which *n_o_* is the largest. The maximum-likelihood tree will instead be the one with the lowest *L_t_,* obtained by minimizing both the left and the right sums in equation 5.

A closer look at one of the simulated datasets where likelihood fails may be helpful in demonstrating this behavior (Supplementary Fig. 2). Two site patterns have a very large impact on the total tree likelihoods of the species tree and the AGT: “11100” (site 1, Fig. 6) and “00110” (site 2, Fig. 6). As expected, the negative log-likelihood (the “cost”) of site pattern 11100 is lower for the species tree than for the AGT (9.91 vs. 11.62; Fig. 6), as this site pattern is concordant with the former and discordant with the latter. Conversely, the cost of site pattern 00110 is lower for the AGT than for the species tree (10.25 vs. 13.11; Fig. 6) because it is concordant with the AGT. Note, however, that the difference in costs of the concordant site patterns in either topology (9.91 - 10.25 = −0.34) is smaller than the difference in costs of discordant sites (13.11 - 11.62 = 1.49). This means that concordant sites cost slightly more on the AGT than the species tree, but that discordant sites cost considerably more on the species tree. Ultimately, this implies that when there are a large number of discordant sites (and the number of concordant sites is not very different between competing topologies; Supplementary Fig. 2), these differences in cost can cause likelihood-based methods to prefer the AGT over the species tree.

Our hypothesis to explain the failure of likelihood methods makes testable predictions. One fundamental prediction is that the length of branches on which sites discordant with one topology and concordant with the other (and vice-versa) are resolved will have a large effect on the final likelihood, and therefore on the tree that is preferred. Importantly, in the topologies considered here these branches are all either tips or an internal branch subtending the entire clade (Fig. 6), and therefore will have no effect on the number of concordant or discordant sites.

We tested this prediction by exploring two more dimensions of tree space: the lengths of branches *w* and z. We ran two new sets of simulations, changing *w* and *z* one at a time. In the first set, *z* was held constant at 1, and *w* was varied from the original value of 12 to either 8 or 20 (dark and light gray shaded regions, respectively; Fig. 7a). The second set of simulations varied the *z* dimension: *w* was held constant at 12, while *z* varied from the original value of 1 to either 0.1 or 10 (dark and light gray shaded regions, respectively; Fig. 7b). These simulations show that, as predicted, the length of branches not directly determining the number of concordant and discordant sites can have a large effect on the region of parameter space in which likelihood fails (parsimony still favors the species tree in all cases; results not shown). Note that non-informative sites and sites discordant with the two competing trees can be resolved along the same branches in both trees; this results in almost identical likelihoods of these sites under the species tree and the AGT (Supplementary Fig. 2; data not shown for non-informative sites). Therefore, although changing the lengths of z and *w* will affect the likelihood of all sites, the impact should be the greatest for sites that are concordant with one tree and discordant with the other, ultimately leading to the different observed maximum-likelihood outcomes between conditions.

**Figure 7:**
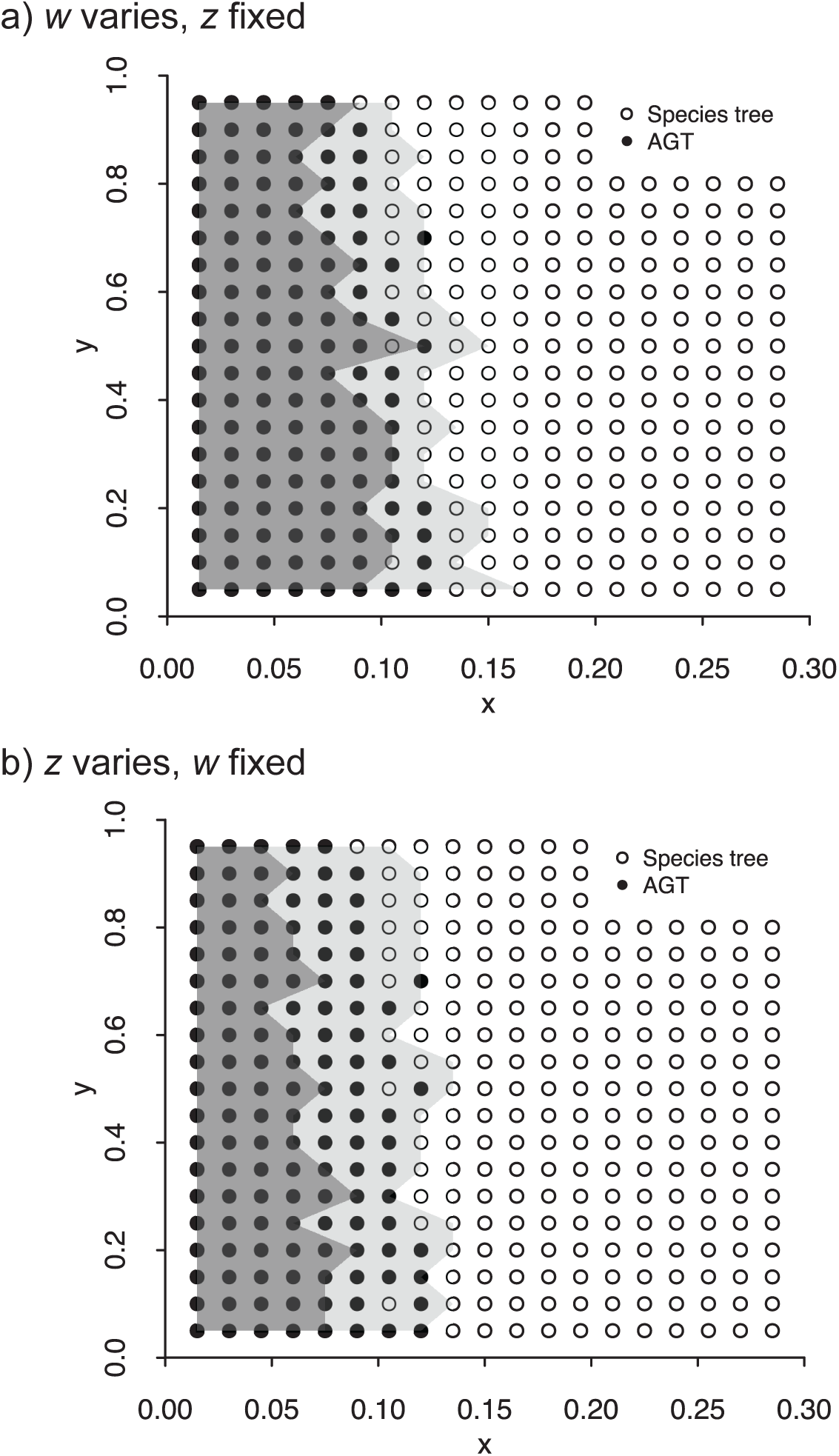
Regions in tree space where likelihood-based methods fail to recover the species tree (black dots and regions shaded in gray). Dots are the same in both (a) and (b), and match those in Fig. 5 (*z* = 1 and *w* = 12 in these simulations). (a) Simulations with *z* = 1, varying *w* to be either *w* = 8 (dark gray) or *w =* 20 (light gray). (b) Simulations with *w =* 12, varying *z* to be either z = 0.01 (dark gray) or z = 10 (light gray).

The results presented in this section are natural mathematical consequences of the theory behind long-established models in phylogenetics. It is likely that they have not been considered before simply because few empirical examples existed in the area of tree space where they become important. But, as discussed more thoroughly below, clarity about the points raised here is critical in avoiding misconceptions about the real cause of the failure of likelihood-based methods, and possibly suggest ways in which these failures can be ameliorated.

## Discussion

The increasing availability of genome-scale datasets has revolutionized and modernized the field of molecular phylogenetics. Sequences from more species and more (longer) genomic segments have provided evolutionary biologists with unprecedented insight into the history of life on Earth. The influx of data has also clarified the relevance and pervasiveness of phenomena that generate gene tree discordance, such as ILS (e.g., Pollard et al. 2006; Hobolth et al. 2011; Brawand et al. 2014; Zhang et al. 2014; Suh et al. 2015; Pease et al. 2016). This in turn has led to the proliferation of methods capable of dealing with discordance (e.g., Liu and Pearl 2007; Than et al. 2008; Liu et al. 2009, 2010; Heled and Drummond 2010; Larget et al. 2010; Mirarab and Warnow 2015; Solís-Lemus and Ané 2016).

When ILS is present at high levels, most gene trees will be discordant with the species tree. Therefore, there has been much interest in the behavior of standard phylogenetic approaches in these instances. Kubatko and Degnan (2007) showed through simulation of a four-taxon phylogeny that the commonly used approach of concatenation and analysis by maximum-likelihood when there are high levels of ILS can lead to strong support for the incorrect tree (the anomalous gene tree; AGT). These authors evaluated the performance of concatenation coupled with likelihood across multiple points of tree space, including in the anomaly zone (Degnan and Rosenberg 2006), in which the topology of the most common gene tree does not match the species tree. Two key conclusions are often drawn from this paper, though neither was one the authors made themselves. The first is that concatenation fails *per se* (i.e., concatenation is the procedure that *directly* causes tree estimation to fail), regardless of the tree-building method used downstream, such as maximum-parsimony or maximum-likelihood (for rare exceptions, see Liu and Edwards 2009; Wu et al. 2014; Degnan and Rhodes 2015; Roch and Warnow 2015; RoyChoudhury et al. 2015; Mirarab et al. 2016). The second conclusion drawn from their study is that the failure of concatenation is caused by the species tree inhabiting the AZ (e.g., Leaché et al. 2015; Olave et al. 2015; Tang et al. 2015; DaCosta and Sorenson 2016; Edwards et al. 2016; Linkem et al. 2016). In fact, both points are addressed directly by Kubatko and Degnan (2007): “Although these results indicate that the existence of an AGT is neither necessary nor sufficient for statistical inconsistency, they demonstrate that [maximum-likelihood] estimation from concatenated sequences can perform poorly for points in or even near the anomaly zone.”

The two apparently common conclusions drawn from early studies noted above are intriguing, but not entirely surprising. While the studies by Kubatko and Degnan (2007) and Liu and Edwards (2009) clearly suggest otherwise, the specific results that speak to these misconceptions might have been obscured by the main findings of these papers, or perhaps by the lack of a clear explanation for the observed inconsistency. Kubatko and Degnan (2007) consider the failure of likelihood-based estimation in certain regions of tree space “surprising,” but do not provide an explanation for it. Liu and Edwards (2009) tentatively suggest that long-branch attraction (Felsenstein 1978) can explain why maximum-parsimony succeeds in recovering the species tree inside the AZ, but do not strongly commit to this hypothesis.

Here, we recapitulate and extend the results of Kubatko and Degnan (2007) and Liu and Edwards (2009) for the same rooted four-taxon species tree, revealing a clear disjunction between the AZ and the region in tree space where likelihood-based estimation fails. Furthermore, we provide a proof for why maximum-parsimony should be successful across this area of tree space under the infinite-site model, and propose a hypothesis to explain why maximum-likelihood trees can be incorrect.

Our results are limited to asymmetric trees in which ILS is observed among four species; that is, when a pair consecutive speciation events (i.e., a pair of nodes) occurs along the same tree path and in a short time span, producing a pectinate topology. Many studies of the AZ have also been limited to four-taxon phylogenies, though some have gone beyond this (e.g., Rosenberg and Tao 2008). While this may seem to limit the generality of the results presented here, we stress that the total number of species in a tree is not the most relevant factor in determining the failure of likelihood-based methods from concatenated data. Instead, what matters is the maximum number of lineages among which ILS occurs, even if only in a small part of a larger tree. Our results imply that as long as the “problematic nodes” of a species tree are limited to four taxa, concatenation coupled with maximum-parsimony or neighbor-joining should succeed in obtaining the correct topology. Extending our analyses to radiations involving five species is tedious, but should be possible. The shape of the anomaly zone for just five species has been shown to be highly complex and to behave in unpredictable ways (Rosenberg and Tao 2008). The exponentially larger numbers of distinct topologies and histories (Degnan and Salter 2005) that result from considering additional species are sure to make the task of predicting the success of parsimony, or of any other method, more difficult. For six taxa (and potentially for more) involved in ILS, it has been shown that maximum-likelihood applied to concatenated data will always result in the wrong species tree (Roch and Steel 2015).

We also note that our demonstration that maximum-parsimony correctly infers the species tree only holds if other phenomena capable of generating phylogenetic incongruence, such as introgression, are not present in the dataset at levels that considerably shift the distribution of gene tree topologies and site patterns. In such cases, the most parsimonious tree is not guaranteed to be correct. Importantly, introgression will also affect most, if not all, other methods that infer species trees (e.g., Leaché et al. 2014; Solís-Lemus et al. 2016). We have also ignored areas of parameter space where parsimony- and distance-based methods will fail for a host of other reasons, including similarity due to homoplasy (e.g., Felsenstein 1978). These problems with parsimony are well known, and nothing presented here should obviate such concerns. However, for datasets using nearly homoplasy-free characters (such as retrotransposon insertions; Suh et al. 2015), our results imply that parsimony can and should be used to avoid problems due to high levels of ILS.

Our results also suggest that the length of the branches leading to and descending from a pair of closely spaced speciation events (denoted *w* and *z* in Fig. 1a) can affect the outcome of maximum-likelihood analyses on concatenated data. These branches are the ones that absorb the cost of discordant site patterns, and as a consequence their length can determine whether the true species tree or an AGT is favored. While many studies vary the lengths of internal branches of a phylogeny to examine the performance of methods for inferring species trees, our results suggest that varying these surrounding branches is necessary to reveal the complete behavior of phylogenetic methods. Our simulations also indicate that if both these branch lengths are short enough, the boundaries of the region in which likelihood-based methods fail will retract, leading to an increased success of these methods in estimating the species tree. This dependence suggests to us that in the future we may find additional ways to ensure the accuracy of maximum-likelihood analyses on concatenated data, possibly by taking into account the substitution rate variation induced by discordance.

## Acknowledgments

We thank Rafael Guerrero for help with the mathematical proof. We also thank Elizabeth Housworth, Laura Kubatko, and David Swofford for comments that helped to improve this work. The research was supported by National Science Foundation grant DEB-1136707.

## Supplementary figure captions

**Supplementary Figure 1:**
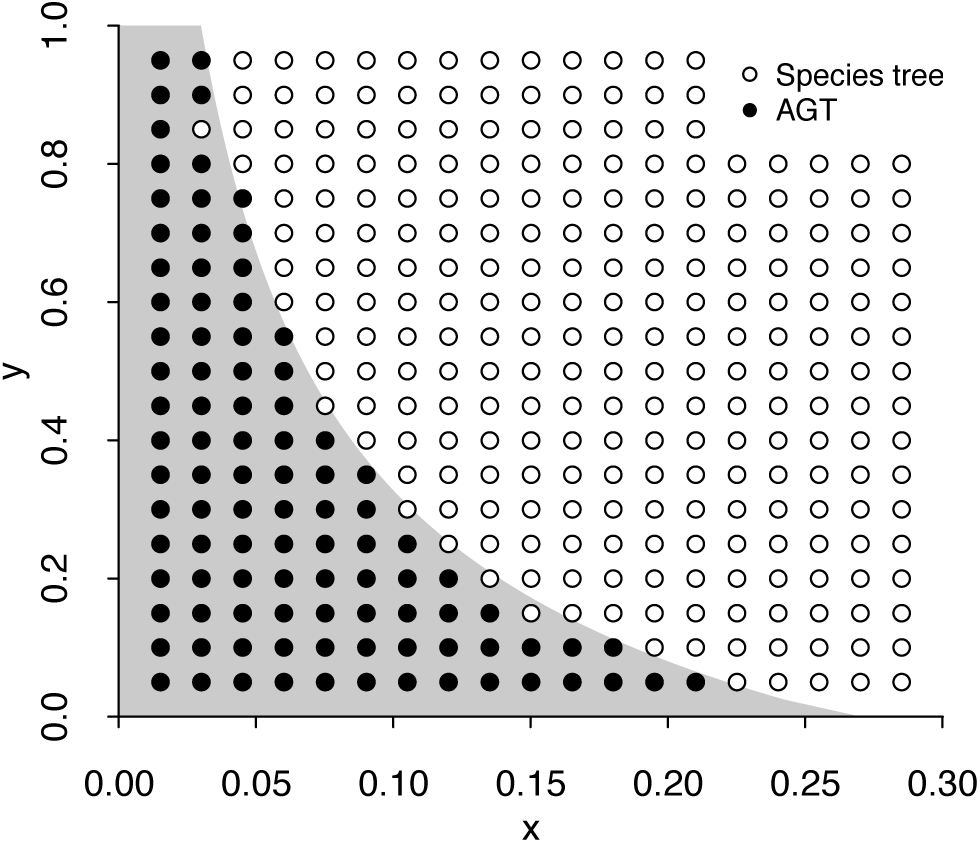
Phylogenetic tree space for species tree ((((A,B),C),D),E), where *x* and *y* correspond to the lengths of the oldest and youngest ingroup internal branches, respectively (as shown in Fig. 1a). The region shaded in gray corresponds to the anomaly zone (derived in Degnan and Rosenberg 2006; and shaded in black in Fig. 1b), in which the most common gene tree is AGT (((A,B),(C,D)),E). Dots show the most common simulated gene tree at each tree space coordinate (out of 20,000 trees).

**Supplementary Figure 2:**
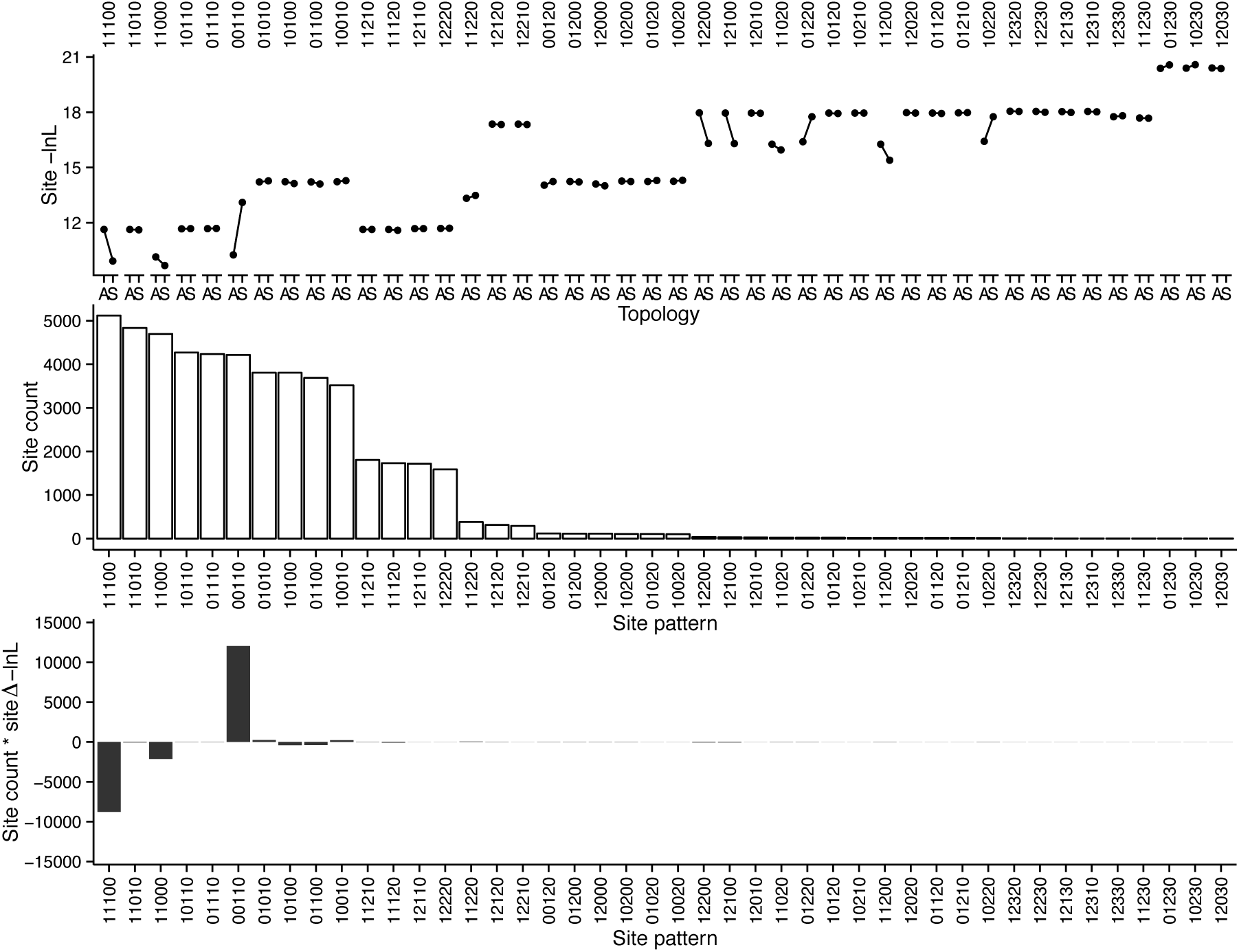
Site likelihoods for the species simulated with *x* = 0.015 and *y* = 0.05. Site patterns are coded based on the different alleles observed at a site (0 corresponds to the nucleotide observed in the outgroup; every other number represents a different, derived nucleotide, regardless of identity). Top panel: the negative log-likelihood (smaller means more likely) of all possible site patterns under the maximum-likelihood branch lengths inferred from a concatenated alignment under the species tree topology (S) or anomalous gene tree topology (A). Middle panel: counts of all site patterns in the concatenated alignment (20,000 1-kb simulated loci; see Appendix B). Bottom panel: the product of the top and middle panels, expressed as the negative log-likelihood difference between the maximum-likelihood trees with the species tree and anomalous gene tree topologies (Δ-lnL = -lnL_s_ - (-lnL_A_)). The species tree is more likely if the likelihood mass above zero summed across all site patterns is greater than that below zero.

## Appendix A

### Calculating S_t_, the overall support for a topology t

For rooted species tree ((A,B),(C,D)) (outgroup omitted), maximum-parsimony methods should recover the topology t that has the largest support (*S_t_*; Eq. A.1 below, but see main text for a more thorough explanation), with support here meaning the total length of gene tree branches that are present as internal branches in topology t. Two topologies compete when data is concatenated: the species tree topology (((A,B),C),D), and the anomalous gene tree (AGT) topology ((A,B),(C,D)). Because these two topologies share the internal branch subtending node {*A,B*}, one can compare *S*_4_ and *S*_1_ (Table 1, main text) by focusing on the branches these two topologies do *not* share: the branch subtending node {*A*, *B*, *C*} (present in the species tree topology) and the branch subtending {C, *D*} (present in the AGT). The species tree topology (*t* = 4; Table 1, main text) will be returned as the most parsimonious (instead of the AGT, *t* = 1) if *S*_4_ >*S*_1_.
*S_t_* is defined in the main text as:

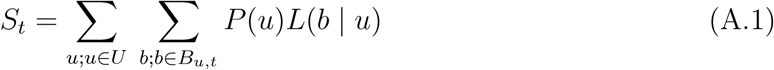

where *U* is the set of gene tree topologies that share internal branches with topology *t*, and *B_u_*_,*t*_ is the set of internal branches that each individual gene tree, *u*, in *U* shares with *t*. *P*(*u*) is the probability of gene tree topology *u* under the species tree (Table 1, main text). L(*b*|*u*) is the expected length in coalescent units (*N*_e_ generations) of branch *b* (in the set *B_u_*,*_t_*) given topology *u*. For the case where the internal branches of the species tree (*x* and *y;* Fig. 1a, main text) have a length of zero (i.e., the species tree is a four-taxon polytomy), finding *L*(*b*|*u*) is straightforward using coalescent theory (Equations 2 and 3, main text).

When the species tree internal branches are not zero, however, a given gene tree topology *u* can be classified into different coalescent history classes (Degnan and Salter, 2005), the set of which is denoted *H*. A history class *h* is defined by the times at which coalescent events take place (Fig. A.1 and Table A.1 and A.2; see below). We can replace the probability of observing each gene tree topology, *P*(*u*), with the probability of each history class *h* in *H* given *u*, *G*(*h* | *u*). Importantly, we must update the definition of *S_t_*, as the expected branch lengths now depend on *h* and *u*:

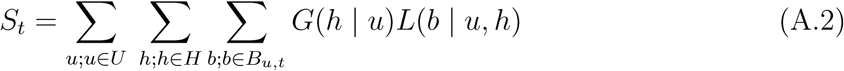

**Figure A.1:**
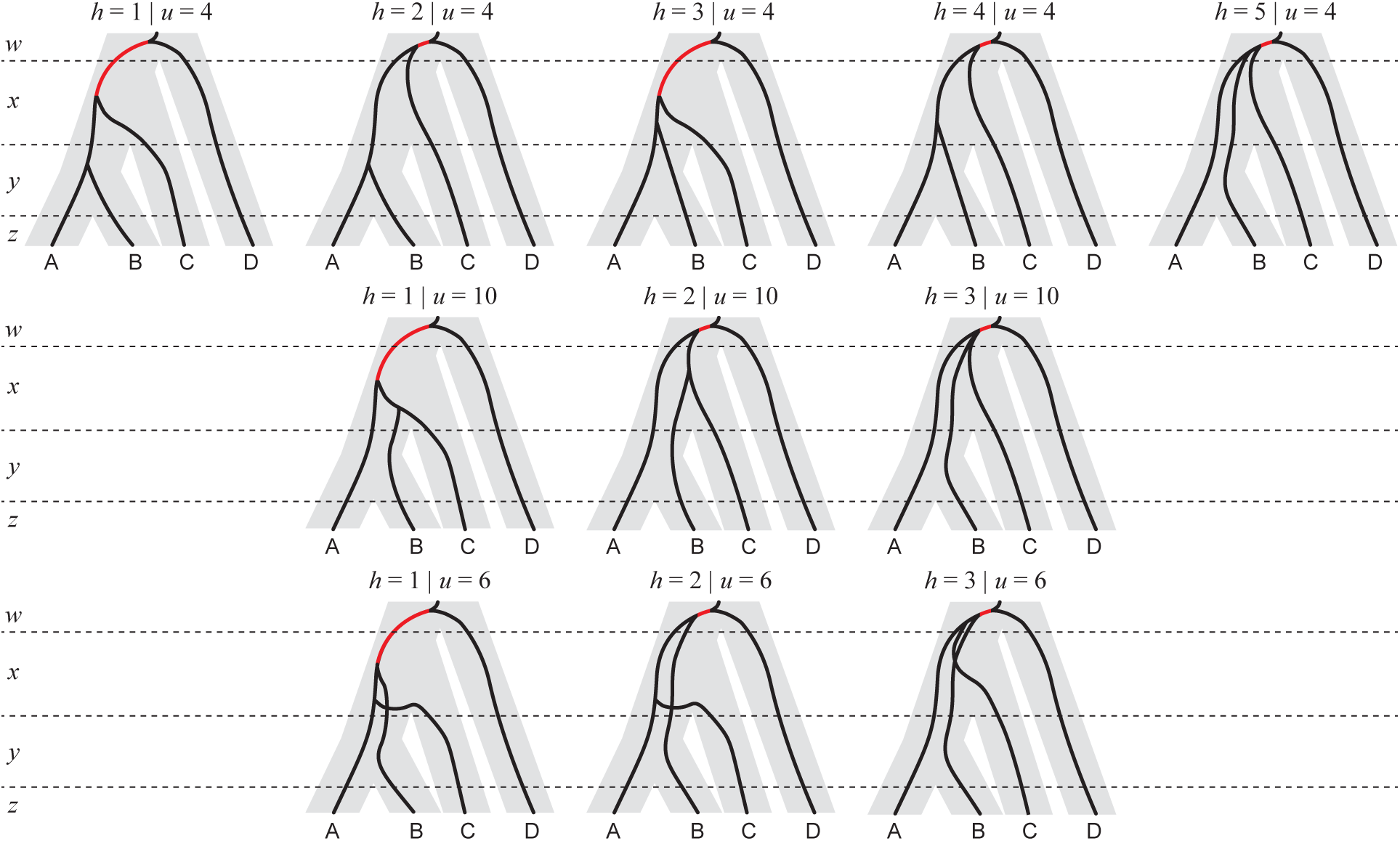
All history classes from all gene tree topologies that share node {A,B,C} with the species tree topology. Branches in red represent the contributed support of each history class to the species tree topology.

**Figure A.2:**
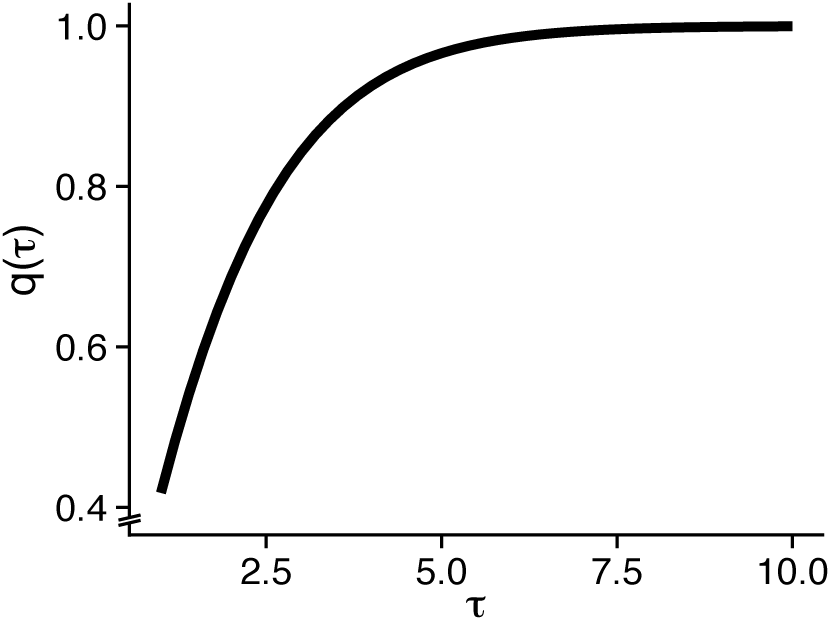
Expected time of coalescence of two lineages within a branch of length *τ*, conditioning on a coalescence event happening.

**Table A.1:** Gene trees supporting the species tree topology through the branch subtending node {A,B,C} (branch lengths in *N*_e_ generations).

| Topology | $u$ | History class, $h$ | Branches containing 1<sup>st</sup> and 2<sup>nd</sup> co-alescences | Probability of history class, $G(h \vert u)$ | Expected branch length, $L(b\vert u, h)$ |
| --- | --- | --- | --- | --- | --- |
| ((AB)C)D) | 4 | 1 | $y, x$ | $g_{21}(y)g_{21}(x)$ | $1 + x - q(x)$ |
| | | 2 | $y, w$ | $\delta_1$ | 1 |
| | | 3 | $x, x$ | $g_{22}(y)g_{31}(x)\frac{1}{3}$ | $1 + x - r(x)$ |
| | | 4 | $x, w$ | $\delta_2$ | 1 |
| | | 5 | $w, w$ | $\delta_3$ | 1 |
| ((BC)A)D) | 10 | 1 | $x, x$ | $g_{22}(y)g_{31}(x)\frac{1}{3}$ | $1 + x - r(x)$ |
| | | 2 | $x, w$ | $\kappa_1$ | 1 |
| | | 3 | $w, w$ | $\kappa_2$ | 1 |
| ((AC)B)D) | 6 | 1 | $x, x$ | $g_{22}(y)g_{31}(x)\frac{1}{3}$ | $1 + x - r(x)$ |
| | | 2 | $x, w$ | $\zeta_1$ | 1 |
| | | 3 | $w, w$ | $\zeta_2$ | 1 |

**Table A.2:** Gene trees supporting the species tree topology through the branch subtending node {C,D} (branch lengths in *N_e_* generations).

| Topology | $u$ | History class, $h$ | Branches containing 1<sup>st</sup> and 2<sup>nd</sup> co-alescences | Probability of history class, $G(h \vert u)$ | Expected branch length, $L(b\vert u, h)$ |
| --- | --- | --- | --- | --- | --- |
| ((AB)(CD)) | 1 | 1 | $y, w$ | $g_{22}(y)g_{33}(x)\frac{1}{3}\frac{1}{3}$ | $1 + \frac{1}{6}$ |
| | | 2 | $x, w$ | $\beta_1$ | 1 |
| | | 3 | $w, w$ | $\beta_2$ | 1 |
| ((CD)A)B) | 14 | 1 | $w, w$ | 1 | $\frac{1}{3}$ |
| ((CD)B)A) | 15 | 1 | $w, w$ | 1 | $\frac{1}{3}$ |

### Calculating the probability of a coalescent history class

The probabilities of coalescent history classes given a gene tree topology (defined here as *G*(*h* | u)) have been derived in Pamilo and Nei (1988) and Rosenberg (2002) for the species tree being considered here (for more general cases, see Degnan and Salter 2005). Those calculations make use of the function *g_ij_*(*τ*) (Tavaré, 1984), defined as:

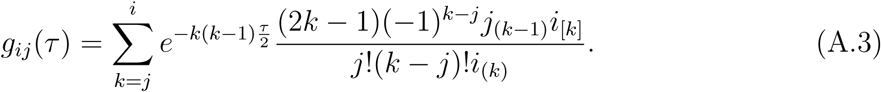

where *a*_(*k*)_=*a*(*a*+1)…(*a*+*k*–1) for *k* ≥ 1 with *a*_(*0*)_=1; and *a*_[*k*]_= *a*(*a*–1)…(*a*–*k*+1) for *k* ≥ 1 with *a*_[0]_ = 1. *g_ij_*(τ) returns the probability that *i* lineages descend from *j* lineages τ coalescent units in the past, with *g_ij_*(τ) = 0 except when *i* ≥ *j* ≥ 1.

From Equation (A.2), comparing *S*_1_ and *S*_4_ requires computing *G(h |* u). Note, however, that because some of the history classes contribute the same support to *S*_t_, we do not have to calculate *G*(*h* | *u*) for all values of *h.* For example, history classes 2, 4 and 5 given *u* = 4 all contribute 1 to *S*_4_, and so their probabilities (*δ*_1_ + *δ*_2_ + *δ*_3_) can be evaluated to 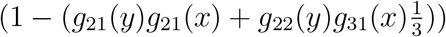 (Table A.1).

### Calculating expected branch lengths

After calculating the probabilities of the different coalescent history classes, *G*(*h* | u), we now must calculate the expected gene tree branch lengths for each *t* contributed by each *h*. For our purposes in comparing the species tree and the AGT, the only branches that matter are those supporting node {*A*, *B, C*} and node {*C, D*}. Evaluating *S*_4_, for example, would entail summing the expected branch lengths in all coalescent histories from all three gene tree topologies that have node {*A*,*B*,*C*} (Fig. A.1; this is equivalent to summing all branches highlighted in red).

Again, expected branch lengths can be obtained with coalescent theory (Tables A.1 and A.2). Some of the expected branch lengths (such as those from history classes 2, 4 and 5, given *u* = 4; Table A.1) are simply the expected time until coalescence of two lineages (*N*_e_ generations = 1 coalescent unit). For the remaining history classes, however, we must find the expected times of coalescence of either two lineages, or three lineages into their MRCA *conditioning* on finding the MRCA within a branch of length *τ*. The former is used when finding the support for the species tree (*t* = 4) coming from history class 1 of the congruent topology (*h* =1 and *u* = 4; Fig. A.1): here, two lineages must coalesce in *x,* so we must subtract the expected time of coalescence (conditioning on it happening in *x*) from 1 + *x*.

In order to derive the expected time of coalescence of two lineages conditioning on a coalescent event happening within a branch of length *τ*, we use the fact that the expected time of coalescence of two lineages, v, is exponentially distributed (with λ = 1), with *pdf*:

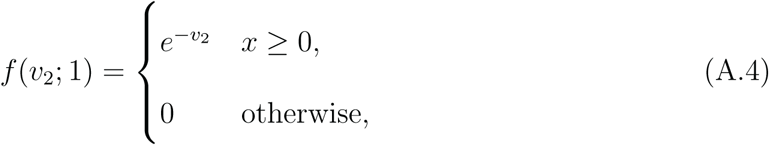

and *cdf:*

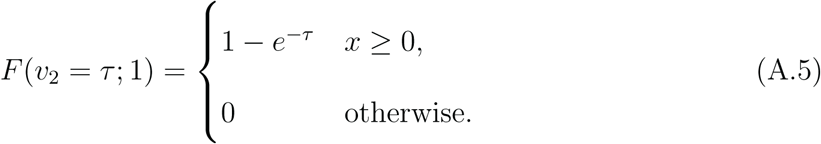

Note that in the *cdf* above, we equate *v*_2_ = *τ* because we are interested in the probability of coalescence before time *τ*.

We can then define the *pdf* of *v*_2_ given that a coalescent event happens within a branch of length *τ*, by dividing Equation (A.4) by Equation (A.5):

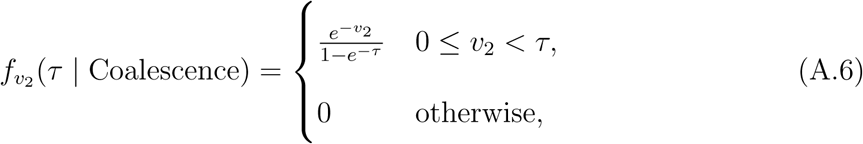

and then finally calculate the *pdf* for the expected time for two lineages to coalesce in a branch of length *τ*, conditioning on a coalescence event happening, *q*(*τ*):

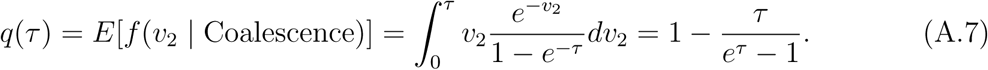

Importantly, *q*(*τ*) converges on 1 coalescent unit, as expected (Fig. A.2).

The same logic outlined above can be used to derive the expected time of coalescence of three lineages into their MRCA within a branch of length τ, conditioning on their coalescence taking place in that branch. In this case, the expected time of coalescence of three lineages into their MRCA, *v*_3_, can be seen as a variable resulting from the convolution of two exponentially distributed random variables (with λ =1 and λ = 3, respectively). If we name the *pdf s* of these two exponential variables *k*(*v*_3_) and *l*(*v*_3_), we can define the *pdf* of the convolved variable:

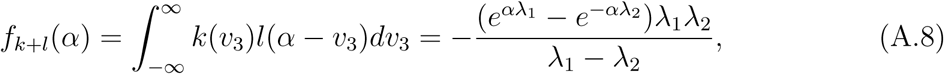

for *α* > 0. Replacing λ_1_ = 1 and λ_2_ = 3, we obtain *pdf*:

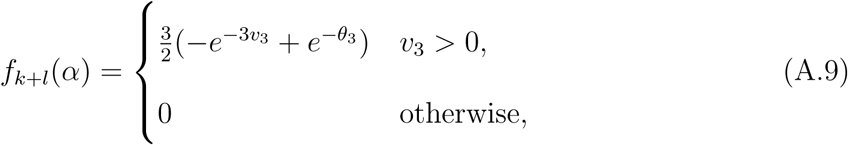

and *cdf* (similarly to what was done above, we equate *v*_3_ = *τ*):

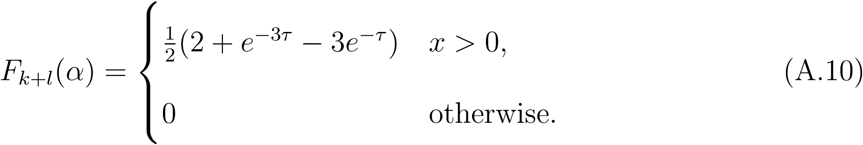

We can then define the *pdf* of *v*_3_ given a coalescent event happens within a branch of length τ, by dividing Equation (A.9) by Equation (A.10):

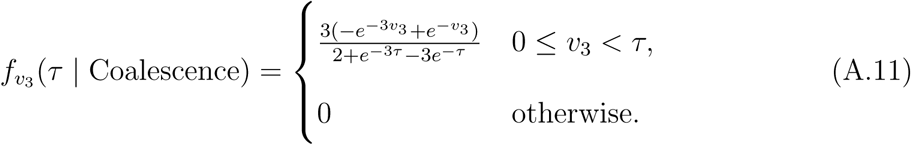

The last step is to calculate the *pdf* for the expected time for two lineages to coalesce in a branch of length τ, conditioning on a coalescence event happening, *r*(*τ*):

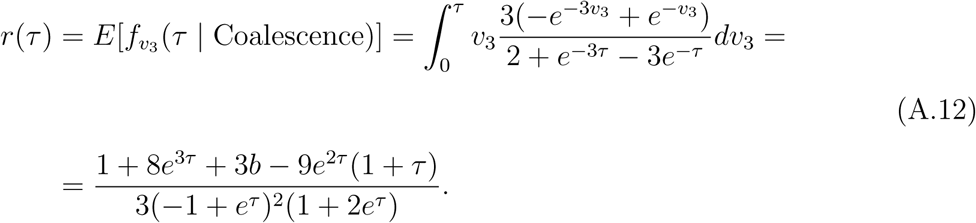

Finally, we must again verify the convergence of *r*(*τ*), except in this case the expectation is 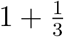 coalescent units (Fig. A.3).

**Figure A.3:**
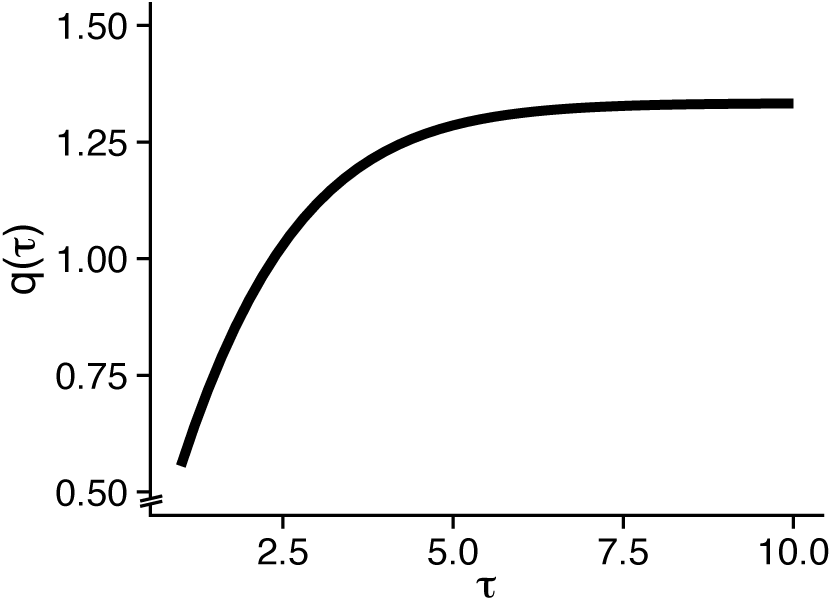
Expected time of coalescence of three lineages within a branch of length *τ*, conditioning on a coalescence event happening.

## Appendix B

### Simulations across the phylogenetic space of a four-taxon species tree

In order to understand the behavior of different tree estimation methods across phylogenetic space, we used the coalescent model to simulate gene trees from an asymmetric species tree with four species in its ingroup, ((((A:*z*,B:*z*):*y*,C):*x*,D):*w*,E), where *z*, *y, x* and *w* are the lengths of terminal branches A and B, and the internal branches subtending (A,B), ((A,B),C) and (((A,B),C),D), respectively. Branch E leads to the outgroup, so the internal branch length *w* was always large enough so no ILS happened between E and any of the remaining taxa.

We explored the phylogenetic space of this species tree by simulating 20,000 gene trees at different x- and y- value combinations (measured in coalescent units, where 1 unit = N_e_ generations), with *x* varying from 0.015 to 0.285 in 0.015 increments, and *y* varying from 0.05 to 0.95 in 0.05 increments – for a total of 361 combinations comprising a square *xy*-grid (*w* and *z* were fixed for this initial set of simulations to 12 and 1 coalescent units, respectively). In addition, we further explored phylogenetic space by simulating along the *xy*-grid four more times: (i) with *z* = 0.1 and *z* =10 (one each; w was fixed at 12 coalescent units), and (ii) with *w* = 8 and *w* = 20 (one each; *z* was fixed at 1 coalescent unit). Simulated gene trees were used in conjunction with the Jukes-Cantor nucleotide evolution model (Jukes and Cantor, 1969) and *θ* = 0.04 to simulate one 1-kb locus alignment per tree. All 20,000 simulated alignments from each *xy*-grid point were concatenated and used in downstream analyses. Coalescent simulations were done with ms (Hudson, 2002) and sequences were simulated with Seq-Gen (Rambaut and Grassly, 1997).

### Comparing empirical and expected support for the species tree and the anomalous tree

We summarized the difference in phylogenetic signal favoring the species tree (SP) versus the anomalous gene tree (AGT) by computing the SP:AGT ratio of the sums of branch lengths supporting each tree. Branch length support for both trees was calculated at 19 grid points along the diagonal of the *xy*-grid (from *x* = 0.015 and y = 0.05, to *x* = 0.285 and *y* = 0.95, and for x = y = 0), with 100 replicates for every point, each replicate consisting of 20,000 gene trees.

For each replicate in each grid point, we computed the support for the species tree by adding the lengths of all internal branches subtending ((A,B),C); these branches were present in 3 of the 15 possible topologies: (((A,B),C),D), (((A,C),B),D), and (((B,C),A),D) (outgroup omitted). Similarly, we added the lengths of all internal branches subtending (C,D) in order to obtain the branch length support for the anomalous tree; these branches are found in topologies ((A,B),(C,D)), (((C,D),A),B), and (((C,D),B),A). Finally, we compared the SP:AGT ratios of branch length support at each grid point to the expected theoretical ratios (see Appendix A).

### Evaluating tree inference methods on concatenated alignments across phylogenetic space

Phylogenies were estimated from the concatenated alignments across the xy-grid using neighbor-joining, parsimony, and maximum-likelihood as implemented in PAUP* v4.0a150 (Swofford, 2002). Maximum-likelihood estimation was done exhaustively, as in Kubatko and Degnan (2007): all 15 possible rooted topologies had their likelihoods evaluated and the top one was reported. We also estimated the maximum-likelihood tree with heuristic search; in this case PAUP* reported one single best tree in all but one point on the grid.

### Inferring site pattern likelihoods under the maximum-likelihood tree

The 20 million sites in each concatenated alignment were first classified into one of 44 unique site pattern bins, after coding the ancestral state (the base present in the outgroup E) as “0”, and the derived states as “1”, “2” or “3” depending on how many different states were present at a given site. This procedure is possible because the Jukes-Cantor model does not incorporate transition-transversion bias, and so site pattern ((((AA)G)G)A), for example, is equivalent to ((((AA)C)C)A); both would be coded as “00110”.

The likelihood of all site patterns was computed for the maximum-likelihood tree at the grid point closest to the origin (*x* = 0.015 and *y* = 0.05). Likelihood computations were done with PAUP*.

## References

Brawand D., Wagner C.E., Li Y.I., Malinsky M., Keller I., Fan S., Simakov O., Ng A.Y., Lim Z.W., Bezault E., Turner-Maier J., Johnson J., Alcazar R., Noh H., Russell P., Aken B., Alföldi J., Amemiya C., Azzouzi N., Baroiller J.-F., Barloy-Hubler F., Berlin A., Bloomquist R., Carleton K., Conte M., D’Cotta H., Eshel O., Gaffney L., Galibert F., Gante H., Gnerre S., Greuter L., Guyon R., Haddad N., Haerty W., Harris R., Hofmann H., Hourlier T., Hulata G., Jaffe D., Lara M., Lee A., MacCallum I., Mwaiko S., Nikaido M., Nishihara H., Ozouf-Costaz C., Penman D., Przybylski D., Rakotomanga M., Renn S., Ribeiro F., Ron M., Salzburger W., Sanchez-Pulido L., Santos M., Searle S., Sharpe T., Swofford R., Tan F., Williams L., Young S., Yin S., Okada N., Kocher T., Miska E., Lander E., Venkatesh B., Fernald R., Meyer A., Ponting C., Streelman J., Lindblad-Toh K., Seehausen O., Palma F. 2014. The genomic substrate for adaptive radiation in African cichlid fish. Nature 513:375–81.

DaCosta J.M., Sorenson M.D. 2016. ddRAD-seq phylogenetics based on nucleotide, indel, and presence-absence polymorphisms: Analyses of two avian genera with contrasting histories. Mol. Phylogenet. Evol. 94:122–154.

Degnan J., Rhodes J. 2015. There are no caterpillars in a wicked forest. Theor. Popul. Biol. 105:17–23.

Degnan J., Rosenberg N. 2006. Discordance of species trees with their most likely gene trees. PLoS Genet. 2:0762–68.

Degnan J.H., Rosenberg N.A. 2009. Gene tree discordance, phylogenetic inference and the multispecies coalescent. Trends Ecol. Evol. 24:332–340.

Degnan J.H., Salter L.A. 2005. Gene tree distributions under the coalescent process. Evolution 59:24–37.

Edwards S., Xi Z., Janke A., Faircloth B., McCormack J., Glenn T., Zhong B., Wu S., Lemmon E., Lemmon A., Leaché A., Liu L., Davis C. 2016. Implementing and testing the multispecies coalescent model: A valuable paradigm for phylogenomics. Mol. Phylogenet. Evol. 94:447–462.

Edwards S.V. 2009. Is a new and general theory of molecular systematics emerging? Evolution 63:1–19.

Felsenstein J. 1978. Cases in which parsimony or compatibility methods will be positively misleading. Syst. Zool. 27:401–410.

Gaut B.S., Lewis P.O. 1995. Success of maximum likelihood phylogeny inference in the four-taxon case. Mol. Biol. Evol. 12:152–162.

Gee H. 2003. Evolution: Ending incongruence. Nature 425:782–782.

Hahn M.W., Nakhleh L. 2016. Irrational exuberance for resolved species trees. Evolution 70:7–17.

Heled J., Drummond A. 2010. Bayesian inference of species trees from multilocus data. Mol. Biol. Evol. 27:570–580.

Hinchliff C., Smith S., Allman J., Burleigh J., Chaudhary R., Coghill L., Crandall K., Deng J., Drew B., Gazis R., Gude K., Hibbett D., Katz L., Laughinghouse H., McTavish E., Midford P., Owen C., Ree R., Rees J., Soltis D., Williams T., Cranston K. 2015. Synthesis of phylogeny and taxonomy into a comprehensive tree of life. Proc. Natl. Acad. Sci. USA 112:12764–12769.

Hobolth A., Dutheil J.Y., Hawks J., Schierup M.H., Mailund T. 2011. Incomplete lineage sorting patterns among human, chimpanzee, and orangutan suggest recent orangutan speciation and widespread selection. Genome Res. 21:349–356.

Huang H., Knowles L. 2009. What is the danger of the anomaly zone for empirical phylogenetics? Syst. Biol. 58:527–536.

Hudson R.R. 1983. Testing the constant-rate neutral allele model with protein sequence data. Evolution 37:203–217.

Huelsenbeck J., Bull J.J., Cunningham C. 1996. Combining data in phylogenetic analysis. Trends Ecol. Evol. 11:152–158.

Huelsenbeck J.P. 1995. Performance of phylogenetic methods in simulation. Syst. Biol. 44:17–48.

Jukes T.H., Cantor C.R. 1969. Evolution of protein molecules. New York: Academic Press p. 21–132.

Kubatko L., Degnan J. 2007. Inconsistency of phylogenetic estimates from concatenated data under coalescence. Syst. Biol. 56:17–24.

Larget B., Kotha S., Dewey C., Ané C. 2010. BUCKy: Gene tree/species tree reconciliation with Bayesian concordance analysis. Bioinformatics 26:2910–2911.

Leaché A., Harris R., Rannala B., Yang Z. 2014. The influence of gene flow on species tree estimation: A simulation study. Syst. Biol. 63:17–30.

Leaché A.D., Chavez A.S., Jones L.N., Grummer J.A., Gottscho A.D., Linkem C.W. 2015. Phylogenomics of Phrynosomatid lizards: Conflicting signals from sequence capture versus restriction site associated DNA sequencing. Genome Biol. Evol. 7:706–719.

Linkem C.W., Minin V.N., Leaché A.D. 2016. Detecting the anomaly zone in species trees and evidence for a misleading signal in higher-level skink phylogeny (Squamata: Scincidae). Syst. Biol. 65:465–477.

Liu L., Edwards S.V. 2009. Phylogenetic analysis in the anomaly zone. Syst. Biol. 58:452–460.

Liu L., Pearl D.K. 2007. Species trees from gene trees: reconstructing Bayesian posterior distributions of a species phylogeny using estimated gene trees distributions. Syst. Biol. 56:504–514.

Liu L., Yu L., Edwards S.V. 2010. A maximum pseudo-likelihood approach for estimating species trees under the coalescent model. BMC Evol. Biol. 10:302.

Liu L., Yu L., Pearl D., Edwards S. 2009. Estimating species phylogenies using coalescence times among sequences. Syst. Biol. 58:468–477.

Mendes F.K., Hahn M.W. 2016. Gene tree discordance causes apparent substitution rate variation. Syst. Biol. 65:711–721.

Mirarab S., Bayzid M., Warnow T. 2016. Evaluating summary methods for multilocus species tree estimation in the presence of incomplete lineage sorting. Syst. Biol. 65:366–380.

Mirarab S., Warnow T. 2015. ASTRAL-II: coalescent-based species tree estimation with many hundreds of taxa and thousands of genes. Bioinformatics 31:i44–i52.

Murphy W.J., Eizirik E., O’Brien S.J., Madsen O., Scally M., Douady C.J., Teeling E., Ryder O.A., Stanhope M.J., de Jong W.W., Springer M.S. 2001. Resolution of early placental mammal radiation using Bayesian phylogenetics. Science 294:2348–2351.

O’Meara B.C. 2012. Evolutionary inferences from phylogenies: A review of methods. Annu. Rev. Ecol. Evol. 43:267–285.

Ogden T., Rosenberg M. 2006. Multiple sequence alignment accuracy and phylogenetic inference. Syst. Biol. 55:314–328.

Olave M., Avila L.J., Sites Jr J.W., Morando M. 2015. Model-based approach to test hard polytomies in the Eulaemus clade of the most diverse South American lizard genus *Liolaemus* (Liolaemini, Squamata). Zool. J. Linn. Soc. 174:169–184.

Page R.D.M. 1996. On consensus, confidence, and “total evidence”. Cladistics 12:83–92.

Pamilo P., Nei M. 1988. Relationships between gene trees and species trees. Mol. Biol. Evol. 5:568–583.

Pease J.B., Haak D.C., Hahn M.W., Moyle L.C. 2016. Phylogenomics reveals three sources of adaptive variation during a rapid radiation. PLoS Biol. 14:e1002379.

Philippe H., Delsuc F., Brinkmann H., Lartillot N. 2005. Phylogenomics. Annu. Rev. Ecol. Evol. 36:541–562.

Pollard D.A., Iyer V.N., Moses A.M., Eisen M.B. 2006. Widespread discordance of gene trees with species tree in *Drosophila:* evidence for incomplete lineage sorting. PLoS Genet. 2:1634–1647.

Robinson D.F., Foulds L.R. 1981. Comparison of phylogenetic trees. Math. Biosci. 53:131–147.

Roch S., Steel M. 2015. Likelihood-based tree reconstruction on a concatenation of aligned sequence data sets can be statistically inconsistent. Theor. Popul. Biol. 100:56–62.

Roch S., Warnow T. 2015. On the robustness to gene tree estimation error (or lack thereof) of coalescent-based species tree methods. Syst. Biol. 64:663–676.

Rokas A., Williams B.L., Carroll S.B. 2003. Genome-scale approaches to resolving incongruence in molecular phylogenies. Nature 425:798–804.

Rosenberg N.A. 2002. The probability of topological concordance of gene trees and species trees. Theor. Popul. Biol. 61:225–247.

Rosenberg N.A., Tao R. 2008. Discordance of species trees with their most likely gene trees: The case of five taxa. Syst. Biol. 57:131–140.

RoyChoudhury A., Willis A., Bunge J. 2015. Consistency of a phylogenetic tree maximum likelihood estimator. J. Stat. Plan. Inference 161:73–80.

Slowinski J.B., Page R.D.M. 1999. How should species phylogenies be inferred from sequence data? Syst. Biol. 48:814–825.

Solís-Lemus C., Ané C. 2016. Inferring phylogenetic networks with maximum pseudolikelihood under incomplete lineage sorting. PLoS Genet. 12: e1005896.

Solís-Lemus C., Yang M., Ané C. 2016. Inconsistency of species tree methods under gene flow. Syst. Biol. 65:843–851.

Soltis P., Soltis D., Chase M. 1999. Angiosperm phylogeny inferred from multiple genes as a tool for comparative biology. Nature 402:402–404.

Steel M., Penny D. 2000. Parsimony, likelihood, and the role of models in molecular phylogenetics. Mol. Biol. Evol. 17:839–850.

Suh A., Smeds L., Ellegren H. 2015. The dynamics of incomplete lineage sorting across the ancient adaptive radiation of neoavian birds. PLoS Biol. 13:e1002224.

Sullivan J., Swofford D.L. 1997. Are guinea pigs rodents? The importance of adequate models in molecular phylogenetics. J. Mamm. Evol. 4:77–86.

Swofford D.L., Waddell P.J., Huelsenbeck J.P., Foster P.G., Lewis P.O., Rogers J.S. 2001. Bias in phylogenetic estimation and its relevance to the choice between parsimony and likelihood methods. Syst. Biol. 500:525–539.

Tajima F. 1983. Evolutionary relationship of DNA sequences in finite populations. Genetics 105:437–460.

Tang L., Zou X., Zhang L., Ge S. 2015. Multilocus species tree analyses resolve the ancient radiation of the subtribe Zizaniinae (Poaceae). Mol. Phylogenet. Evol. 84:232–239.

Than C., Ruths D., Nakhleh L. 2008. PhyloNet: a software package for analyzing and reconstructing reticulate evolutionary relationships. BMC Bioinformatics 9:322.

Tonini J., Moore A., Stern D., Shcheglovitova M., Ortí G. 2015. Concatenation and species tree methods exhibit statistically indistinguishable accuracy under a range of simulated conditions. PLoS Curr. 7.

White M.A., Ané C., Dewey C.N., Larget B.R., Payseur B.A. 2009. Fine-scale phylogenetic discordance across the house mouse genome. PLoS Genet. 5:e1000729

Wu Y.-C., Rasmussen M.D., Bansal M.S., Kellis M. 2014. Most parsimonious reconciliation in the presence of gene duplication, loss, and deep coalescence using labeled coalescent trees. Genome Res. 24:475–486.

Zhang G., Li C., Li Q., Li B., Larkin D.M., Lee C., Storz J.F., Antunes A., Greenwold M.J., Meredith R.W., Ödeen A., Cui J., Zhou Q., Xu L., Pan H., Wang Z., Jin L., Zhang P., Hu H., Yang W., Hu J., Xiao J., Yang Z., Liu Y., Xie Q., Yu H., Lian J., Wen P., Zhang F., Li H., Zeng Y., Xiong Z., Liu S., Zhou L., Huang Z., An N., Wang J., Zheng Q., Xiong Y., Wang G., Wang B., Wang J., Fan Y., Fonseca R., Alfaro-Núñez A., Schubert M., Orlando L., Mourier T., Howard J., Ganapathy G., Pfenning A., Whitney O., Rivas M., Hara E., Smith J., Farré M., Narayan J., Slavov G., Romanov M., Borges R., Machado J., Khan I., Springer M., Gatesy J., Hoffmann F., Opazo J., Håstad O., Sawyer R., Kim H., Kim K.-W., Kim H., Cho S., Li N., Huang Y., Bruford M., Zhan X., Dixon A., Bertelsen M., Derryberry E., Warren W., Wilson R., Li S., Ray D., Green R., O’Brien S., Griffin D., Johnson W., Haussler D., Ryder O., Willerslev E., Graves G., Alström P., Fjeldså J., Mindell D., Edwards S., Braun E., Rahbek C., Burt D., Houde P., Zhang Y., Yang H., Wang J., Jarvis E., Gilbert M., Wang J., Ye C., Liang S., Yan Z., Zepeda M., Campos P., Velazquez A., Samaniego J., Avila-Arcos M., Martin M., Barnett R., Ribeiro A., Mello C., Lovell P., Almeida D., Maldonado E., Pereira J., Sunagar K., Philip S., Dominguez-Bello M., Bunce M., Lambert D., Brumfield R., Sheldon F., Holmes E., Gardner P., Steeves T., Stadler P., Burge S., Lyons E., Smith J., McCarthy F., Pitel F., Rhoads D., Froman D. 2014. Comparative genomics reveals insights into avian genome evolution and adaptation. Science 346:1311–1320.

Zuckerkandl E., Pauling L. 1965. Molecules as documents of evolutionary history. J. Theor. Biol. 8:357–366.

## References

Degnan, J. H. and L. A. Salter. 2005. Gene tree distributions under the coalescent process. Evolution 59:24–37.

Hudson, R. R. 2002. Generating samples under a wright-fisher neutral model. Bioinformatics 18:337–338.

Jukes, T. H. and C. R. Cantor. 1969. Evolution of protein molecules. Academic press, New York.

Kubatko, L. S. and J. H. Degnan. 2007. Inconsistency of phylogenetic estimates from concatenated data under the coalescence. Systematic Biology 56:17–24.

Pamilo, P. and M. Nei. 1988. Relationships between gene trees and species trees. Molecular Biology and Evolution 5:568–583.

Rambaut, A. and N. C. Grassly. 1997. Seq-Gen: an application for the Monte Carlo simulation of DNA sequence evolution along phylogenetic trees. Computer Applications in the Biosciences 13:235–238.

Rosenberg, N. A. 2002. The probability of topological concordance of gene trees and species trees. Theoretical Population Biology 61:225–247.

Swofford, D. L. 2002. PAUP*. Phylogenetic analysis using parsimony (*and other methods). Version 4. Sinauer Associates, Sunderland, MA.

Tavaré, S. 1984. Line-of-descent and genealogical processes, and their application in population genetics models. Theoretical Population Biology 26:119–164.

